# Reconstitution of prospermatogonial specification *in vitro* from human induced pluripotent stem cells

**DOI:** 10.1101/2020.09.08.286930

**Authors:** Young Sun Hwang, Shinnosuke Suzuki, Yasunari Seita, Jumpei Ito, Yuka Sakata, Hirofumi Aso, Kei Sato, Brian P. Hermann, Kotaro Sasaki

**Author notes:** Correspondence should be addressed to: Kotaro Sasaki, M.D., Ph.D. These authors contributed equally to this work (Young Sun Hwang, Shinnosuke Suzuki, Yasunari Seita).

## Abstract

Establishment of spermatogonia throughout the fetal and postnatal period is essential for production of spermatozoa and male fertility. Here, we established a protocol for *in vitro* reconstitution of human prospermatogonial specification whereby human primordial germ cell (PGC)-like cells (hPGCLCs) differentiated from human induced pluripotent stem cells were further induced into M-prospermatogonia-like cells (MLCs) and T1 prospermatogonia-like cells (T1LCs) using long-term cultured xenogeneic reconstituted testes. Single cell RNA-sequencing was used to delineate the lineage trajectory leading to T1LCs, which closely resemble human T1-prospermatogonia *in vivo* and exhibited gene expression related to spermatogenesis and diminished proliferation, a hallmark of quiescent T1 prospermatogonia. Notably, this system enabled us to visualize the dynamic and stage-specific regulation of transposable elements during human prospermatogonial specification. Together, our findings pave the way for understanding and reconstructing human male germline development *in vitro*.

## INTRODUCTION

Germ cells (GCs) are the only cells that transmit genetic information from one generation to the next. Moreover, the germline lineage is the foundation for totipotency, enabling the development of all cells and tissues within an individual organism. The germline is established as primordial germ cells (PGCs), undergo a complex cascade of developmental processes, resulting in the formation of either spermatozoa or oocytes depending on the sex-specific niche signals provided by the gonad^1^ . Accordingly, errors that occur during any of these steps can lead to various critical conditions, including infertility or congenital anomalies. Therefore, a precise understanding of the mechanisms regulating human GC development has significant implications, not only for biology in general, but also for a broad range of human diseases.

Males are responsible for nearly 50% of infertility among reproductive-age couples. Non-obstructive azoospermia (NOA) characterized by the lack of sperm in ejaculate affects about 10% of such men, the etiology of which remains unknown in nearly half^2–5^. The most severe form of NOA, Sertoli cell only syndrome (SCOS), is characterized by complete lack of male GCs and likely arises due to disruption of male GC development in the fetal and/or early postnatal stages^6–8^. In mice, male-specific GC development starts when PGCs colonize fetal testis, and immediately differentiate into prospermatogonia (also known as gonocytes) which divide several times as M-prospermatogonia and then become mitotically quiescent as T1-prospermatogonia (T1). Immediately after birth, T1, centrifugally migrate from the center of seminiferous cords towards the periphery, the site of the spermatogonial stem cell (SSC) niche^9–12^. Concomitantly, T1 resume proliferation and differentiate into T2-prospermatogonia (T2), which are considered the immediate precursors to spermatogonial stem cells (SSCs), a founding population to establish ongoing spermatogenesis^9, 10^. Historically, male GCs at the prespermatogenesis (gonadal) phase have been referred to under various terms including gonocytes, fetal spermatogonia, or prespermatogonia^10^. In this paper, we employed the above nomenclature (M-, T1-, and T2-prospermatogonia) originally proposed by Hilscher et al., which clearly conveys GC identity in both sex and timing of development^10, 13^. We also refer to all pregonadal phase (migratory and pre-migratory) GCs as PGCs.

Using single cell RNA-seq on human fetal testicular cells, Li et al. demonstrated two distinct GC types, mitotic and mitotic-arrest human fetal germ cells (FGCs)^14^, which by definition, appear to correspond to M and T1 in mice. Interestingly, in contrast to mice, the transition of M into T1 occurs asynchronously during the first and second trimester^14^. Moreover, unlike murine T1 that predominantly localize within the tubular lumen, at least some of the M and T1 in humans and non-human primates already reside on the basement membranes of seminiferous cords^14–16^. Such spatiotemporal heterogeneity is unique to primates and suggests species-specific divergence of prospermatogonial specification.

Given such divergence, approaches to directly address the mechanisms of human male GC development are required to further our understanding of this process and provide more relevant and novel insights into the causes of human male infertility. Such an objective, however, has been hampered by both ethical and technical constraints in interrogating human fetal GCs, which are scarce and do not persist into adulthood. A reliable *in vitro* reconstitution method that accurately recapitulates fetal phases of human male GC development will allow scalable expansion, manipulation and visualization of developing male GCs, thereby providing the ability to investigate the molecular mechanisms of human male GC development. To this end, we previously established robust *in vitro* methods to induce human inducible pluripotent stem cells (hiPSCs) into human PGC-like cells (hPGCLCs), which resemble pre-migratory stage PGCs^17^. More recently, successful maturation of female hPGCLCs into pre-meiotic oogonia-like cells using xenogeneic reconstituted ovaries (xrOvaries) has also been reported^18^. However, approaches permitting hPGCLCs differentiation into prospermatogonia and beyond have not been reported. In this study, we established an analogous culture method, which we termed the xenogeneic reconstituted testis (xrTestis) culture, which allows for the differentiation of hPGCLCs into T1LCs. Notably, T1LCs generated under these conditions bear a transcriptome that closely resembles the transcriptome of T1 *in vivo*, which we defined using single cell profiling of testicular samples at 17-18 week gestational age. Together, this culture method recapitulates *in vivo* human male GC development, and allows us to understand the genetic pathways governing human male GC development.

## RESULTS

### Characterization of prospermatogonia in human fetal testes at 2^nd^ trimester

Precise understanding of the lineage trajectory of male GCs *in vivo* is a prerequisite for validating human male GC development reconstituted *in vitro*. During the 2^nd^ trimester, human fetal testes consist of heterogenous cell types at different developmental stages, including the proliferative M and T1 in mitotic arrest^14, 19^. Therefore, we set out to profile human fetal testes obtained from 3 donors with gestational age of 17W3d (Hs31), 18W0d (Hs26), 18W5d (Hs27) (Supplementary Fig. 1a-e). Histologic sections revealed compact seminiferous cords with a cylindrical shape embedded in highly cellular stroma (Supplementary Fig. 1a). Seminiferous cords showed scattered GCs with large vesicular nuclei and prominent nucleoli. Stroma contained many cells with round nuclei with abundant eosinophilic cytoplasm, features characteristic of fetal Leydig cells (FLCs)^16^. To determine the linage trajectory of these cells, we next performed single cell RNA-seq (scRNA-seq) using a 10x Genomics platform. Cell suspension of Hs26 and Hs27 testes were cryopreserved previously and Hs31 was freshly prepared before use in single cell analyses. Out of a total of ∼18,000 cells for which transcriptomes were available, 16,429 cells remained for downstream analysis after filtering out low quality cells. Among these, we detected ∼1900-2700 median genes/cell at a sequencing depth of 22-69k mean reads/cell (Supplementary Fig. 1b). By profiling the expression of known marker genes in a tSNE plot^14, 20–23^, we identified clusters representing multiple known cell types of fetal testis including *DND1*+ GCs, *SOX9*+ Sertoli cells (SCs), *INSL3*^+^/*CYP17A1*^+^ fetal Leydig cells (FLCs), *TCF21*^+^ stromal cells [ST, also described as Leydig precursors^14^] and *KDR*+ endothelial cells (ECs) (Supplementary Fig. 1c, d). We also identified minor cell types that were relatively uncharacterized previously in human fetal testis, including *HLA*-*DRB1*^+^ macrophages (MΦ) and *MYH11*^+^ smooth muscle cells (SMCs)^24, 25^. To identify marker genes characteristic of annotated clusters, we performed differentially expressed gene (DEG) analysis, comparing each cluster with remaining clusters (Supplementary Fig. 1e, Supplementary Table 1). The number of DEGs was highest in GCs, consistent with their unique biological characteristics. GC-specific DEGs were enriched in genes bearing GO terms such as “DNA repair” and “spermatogenesis” (Supplementary Table 1). DEGs in Leydig cells were enriched in genes for “cholesterol biosynthetic process” or “steroid biosynthetic process”, suggesting that these cells may produce androgens ^26^. DEGs in ST were enriched for terms such as “extracellular matrix” or “collagen catabolic process”, which may indicate their role in scaffolding testicular tissue architecture (Supplementary Table 1). Additional key markers identified by DEG analysis and GO terms enriched for DEGs were shown in Supplementary Fig. 1d, e and Supplementary Table 1.

We next performed re-clustering analysis focusing only on the GCs, which revealed two distinct clusters (Fig. 1a). We found that a cluster at the upper right portion of the tSNE plot expresses known markers for M (mitotic FGCs) and PGCs (migrating FGCs), such as *POU5F1*, *NANOS3*, *TFAP2C* (Fig. 1b)^14, 27, 28^. Another cluster at the lower left portion of the plot expressed markers for T1 (mitotic-arrest FGCs) such as *PIWIL4*, *TEX15* or *RHOXF1*^14, 29^. *DDX4* expression was also upregulated in this cluster consistent with previous IF studies using it as a marker for human T1^14, 30^ although weaker expression was also seen in M (Fig. 1b). Accordingly, IF studies showed two population within seminiferous cords, POU5F1^+^DDX4^+^ (388/853, 45.5%) and POU5F1^-^DDX4^+/++^ (465/853, 54.5%) cells, representing M or T1, respectively. T1 exhibit significantly lower transcript levels for proliferation markers, such as *AURKB*, *CCNA2*, *MKI67*, and *TOP2A* and marked reduction of MKI67 protein expression by IF (Fig. 1c, d, e)^31^, confirming that these T1 are indeed at the mitotic-arrest stage. RNA velocity analysis using nascent transcripts further corroborated the overall lineage trajectory from M to T1 (Supplementary Fig. 1f)^32^.

**Fig. 1.**
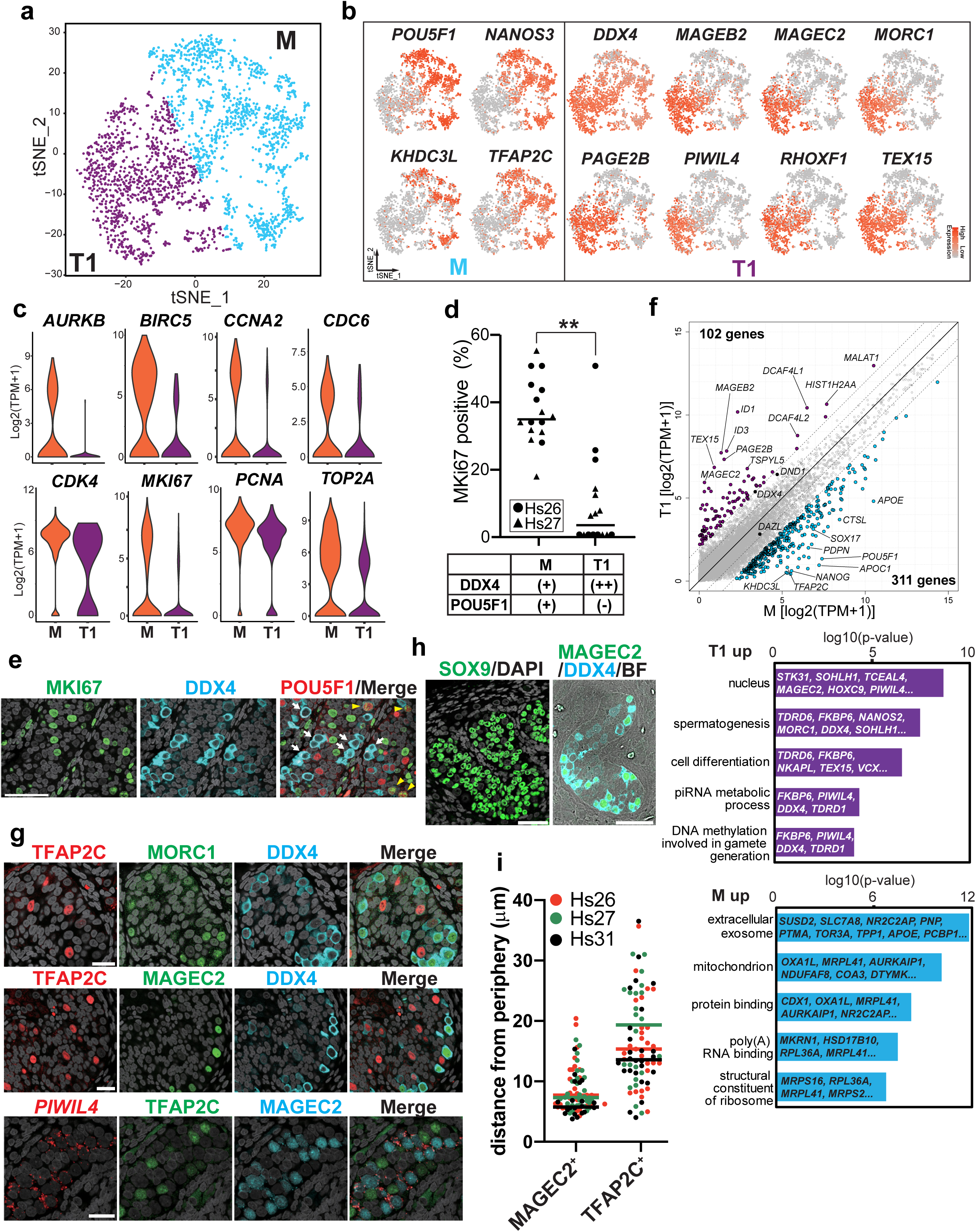
Characterization of germ cells in human fetal testis at 2^nd^ trimester. **a** tSNE plot focused on germ cells (GCs) defined in Fig. S1C. Cluster identities are determined by expression of markers projected on the same tSNE plot in (**b**). M, M-prospermatogonia; T1, T1-prospermatogonia. **b** tSNE feature plots of known and newly identified markers for M or T1. **c** Violin plots of proliferation markers for M (1419 cells) or T1 (1170 cells) defined in (**a**) and (**b**). FDR < 10^-65^ for all markers. **d** Percentages of MKI67^+^ cells among DDX4^+^POU5F1^+^ M or DDX4^++^POU5F1^-^ T1 present in Hs26 (18W0d, black circle) or Hs27 (18W5d, black triangle) samples as assessed by IF. Each dot represents the count per tubular cross section. **e** IF images of a paraffin section of fetal testis (Hs27) for MKI67 (green) and DDX4 (cyan), merged with DAPI (white) and/or POU5F1(red). White arrows indicate DDX4^++^POU5F1^-^ T1 and yellow arrowheads indicate DDX4^+^POU5F1^+^ M co-expressing MKI67. Scale bar, 50 μm. **f** Scatter plot for comparison of averaged gene expression levels between M and T1 (top). Cyan, genes higher in M; Purple, genes higher in T1 (more than 4-fold difference [flanking diagonal lines], mean long2[TPM+1] >2, FDR <0.01). Key genes are annotated and the number of DEGs are indicated. GO analysis of the DEGs with p-value and representative genes contributing to each GO term are shown (bottom). **g** IF images of paraffin sections of Hs31 for TFAP2C (red), DDX4 (cyan) combined with MORC1 (green, top) or MAGEC2 (green, middle) and images for combined *in situ* hybridization (co-ISH) for *PIWIL4* (red) with IF for TFAP2C (green) and MAGEC2 (cyan) (bottom). All images are merged with DAPI (white). Merged images for all 4 color channels are shown at far right. Bar, 25 μm. **h** IF images of paraffin sections of Hs26 for SOX9 (green) merged with DAPI (left) or for MAGEC2 (green) and DDX4 (cyan) merged with bright field (BF) (right). IF for SOX9 and BF highlight the border between tubules and the stroma. Bar, 50 μm. **i** Distances (μm) from the periphery of tubules for TFAP2C^+^MAGEC2^-^ M or TFAP2C^-^ MAGEC2^+^ T1 as quantified by IF images for Hs26 (red), 27 (green) and 31 (purple). Bars indicate median value for each cell type per sample. See also Supplementary Fig. 1 and Supplementary Table 1.

Pairwise DEGs between T1 and M revealed sharp downregulation of GC specifier genes (*SOX17, TFAP2C, PRDM1, SOX15, NANOS3*) and pluripotency-associated genes (*POU5F1, NANOG, TCL1B, TFCP2L1*) as M differentiate into T1 (Fig. 1b. f, Supplementary Table 2). SOX2 is not expressed in either cell types^27, 28, 33^. 311 DEGs identified in M were enriched for those bearing GO terms such as “mitochondrial inner membrane” and “mitochondrial respiratory chain complex I assembly”, suggesting that oxidative phosphorylation may be activated in M as reported in mice, and that this process might be turned off as cells differentiate into T1^34^. 102 DEGs upregulated during the M-to-T1 transition include X-linked cancer-testis antigens belonging to the MAGE and PAGE gene families, such as *MAGEA4, MAGEB2, MAGEC2, PAGE1, PAGE2,* and *PAGE2B*. Many genes previously recognized to mark prospermatogonia (*RHOXF1, NANOS2, DDX4*) or adult spermatogonia (*SIX1, DCAF4L1, PLPPR3, EGR4*) were also among these DEGs^14, 29, 35^. Accordingly, GO terms in these DEGs included “spermatogenesis” (Fig. 1f). Notably, the majority of genes involved in piRNA pathways (e.g. *PIWIL4, TEX15, MORC1*), key guardians of genomic integrity during spermatogenesis, were highly upregulated in T1 in concordance with GO terms, such as “piRNA metabolic process” or “DNA methylation involved in gamete generation” (Fig. 1f). Some of the markers listed among the top DEGs, such as *MAGEC2, MORC1* were not previously recognized as markers of human T1. IF analysis revealed discrete nuclear immunoreactivity for MORC1 and MAGEC2 only in peripherally located DDX4^+^ T1 (Fig. 1g-i). On the other hand, TFAP2C exclusively marked centrally located M. In-situ hybridization (ISH) analysis showed that signals for *PIWIL4* was specifically localized at the perinuclear regions of MAGEC2^+^ T (Fig. 1g). Overall, these findings clearly delineated two distinct male GC types, M and T1 harboring unique characteristics in human fetal testes.

### Establishment of male hiPSCs bearing the TFAP2C-2A-EGFP (AG); DDX4/hVH-2A-tdTomato (VT); PIWIL4-ECFP (PC) alleles (9A13 AGVTPC)

Having defined the transcriptomics of prospermatogonial development at high resolution, we embarked on the *in vitro* reconstitution of this process using hiPSCs as starting material.

Our transcriptomic analysis, coupled with previous reports in humans and non-human primates indicate that *DDX4* and *PIWIL4* expression mark T1, and that the expression of both genes is maintained at least until spermatogenesis commences^14, 15, 36^. *DDX4* expression is likely upregulated earlier than *PIWIL4* given the weaker but significant levels of expression of *DDX4* at M (Fig. 1b)^14^. Moreover, *TFAP2C*, a marker for PGCs, was swiftly downregulated upon differentiation into T1 (Fig. 1b, f, g). These observations led us to hypothesize that a combination of *TFAP2C*, *DDX4* and *PIWIL4* would serve as a powerful marker to visualize the transition from hPGCLCs into the prospermatogonial stage. To this end, we introduced targeted *DDX4/hVH-2A-tdTomato* (VT) and *PIWIL4-ECFP* (PC) alleles into the previously established *TFAP2C-2A-EGFP* (AG) hiPSCs (585B1 1-7, XY)^17^ to generate hiPSCs bearing triple knock-in fluorescence reporters (AGVTPC) (Supplementary Fig. 2a-g). We identified one clone, 9A13, harboring biallelic targeting of both VT and PC (Supplementary Fig. 2c, d). 9A13 hiPSCs could be stably maintained under feeder free condition and exhibited a normal male karyotype (46, XY) (Supplementary Fig. 2e). They formed round and tightly packed colonies, characteristic of hiPSCs (Supplementary Fig. 2f) and expressed pluripotency-associated genes, POU5F1, SOX2 and NANOG (Supplementary Fig. 2g). We also confirmed that 9A13 hiPSCs were able to differentiate into hPGCLCs through incipient mesoderm-like cells (iMeLCs) with an induction efficiency of ∼53% of AG^+^ hPGCLCs (Supplementary Fig.2h, i, j, k), consistent with a previous study^17^.

### Establishment of xenogeneic reconstituted testis (xrTestis)

A previous study successfully reconstituted mouse fetal prospermatogonia from mESC-derived PGC-like cells (mPGCLCs) using reconstituted testes, in which dissociated mouse fetal testicular somatic cells were mixed with mPGCLCs before culture^37^. As a first step in applying this methodology to humans, we examined whether dissociated cells from mouse fetal testes could be reassembled in the absence of mouse(m)PGCs or mPGCLCs (Fig. 2a).

**Fig. 2.**
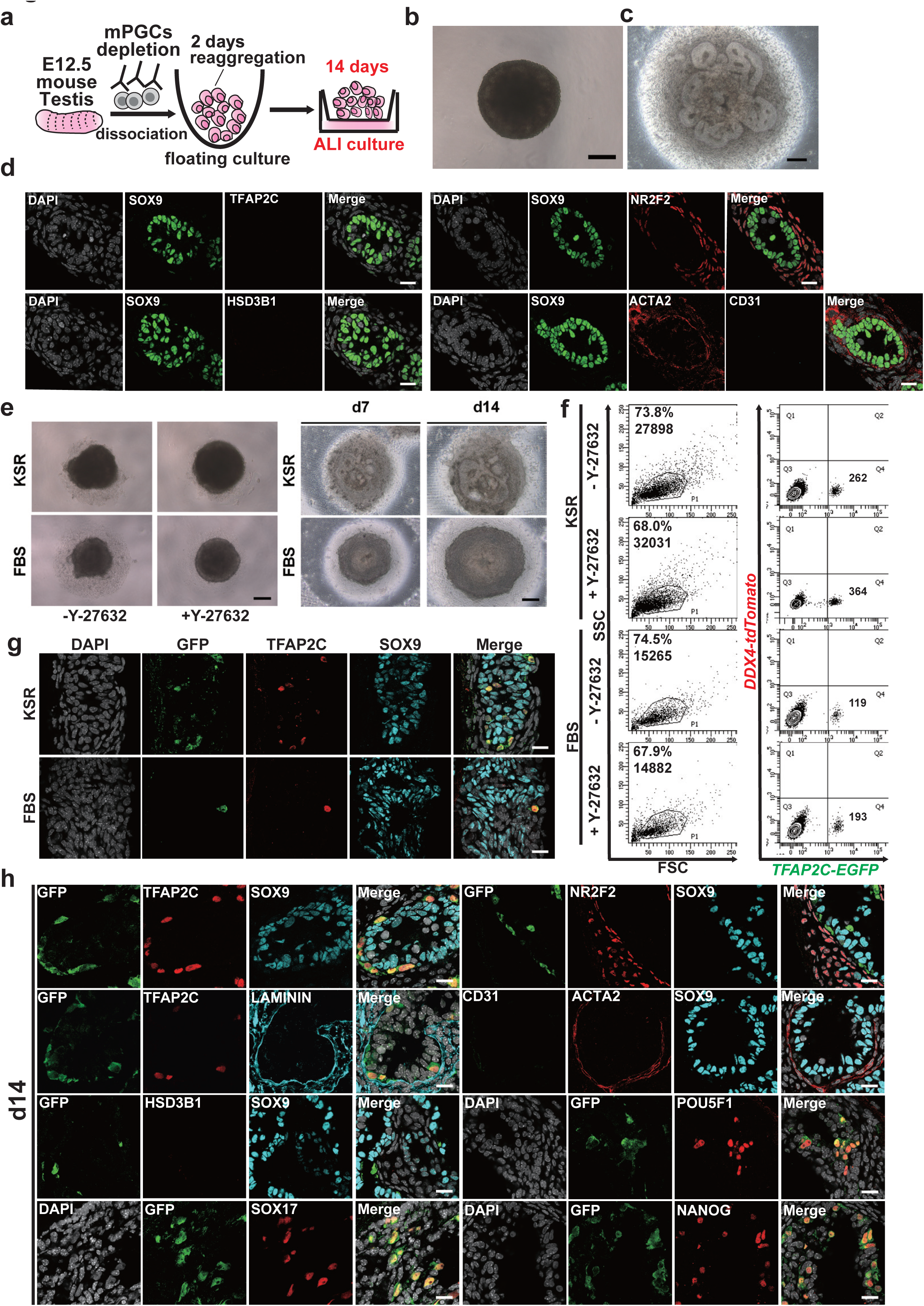
Optimization of rTestis and xrTestis culture. **a** Scheme for rTestis culture using mouse testicular somatic cells at embryonic day (E) 12.5. Mouse PGCs (mPGCs) are depleted by MACS. ALI, air-liquid interphase; rTestis, reconstituted testis. **b, c** BF images of d2 floating aggregate (**b**) and d14 xrTestis on ALI culture (**c**). Bars, 200 µm. **d** IF images of rTestes at d14 for GC (red: TFAP2C) and somatic cell markers (green: SOX9; red: NR2F2, HSD3B1, ACTA2; cyan: CD31) with merges with DAPI (white). Bar, 20 µm. **e** BF images of d2 floating aggregates (left), and d7 and d14 xrTestes (right) cultured in KSR-based or FBS-based medium. Floating aggregates are cultured in the presence or absence of Y-27632 (left). Y-27632 is included in all xrTestis culture (right). Bars, 200 µm. **f** FACS analysis of d2 floating aggregates cultured in KSR or FBS-based medium to assess the number of total cells [in dot plot showing SSC (side scatter) and FSC (forward scatter), left] and the number of hPGCLC-derived cells (TFAP2C-EGFP [AG]-positive, right). The percentages of cells in P1 gates (living cells), the total cell numbers (left) and the numbers of AG-positive cells per floating aggregate (right) are shown. **g** IF images of d14 xrTestes cultured in KSR- or FBS-based medium for GFP (green), TFAP2C (red), SOX9 (cyan) and DAPI (white) with their merges. Bar, 20 µm. **h** IF images of d14 xrTestes for GC markers (red: TFAP2C, POU5F1, SOX17 or NANOG), a marker for hPGCLC-derived cells (green: GFP), a basement membrane marker (cyan: LAMININ), somatic cell markers (green: CD31; red: NR2F2, ACTA2, HSD3B1; cyan: SOX9) and DAPI (white) with their merges. Bars, 20 µm. See also Supplementary Fig. 2 and 3.

E12.5 mouse fetal testicular cells depleted of mPGCs readily formed tight aggregates upon floating culture for 2 days (reconstituted testis, rTestis) (Fig. 2b). Upon an additional 14 days of culture on an air-liquid interface (ALI), rTestes exhibited numerous anastomosing tubular structures observable with brightfield microscopy and were confirmed to consist of tubular structures comprised of SOX9^+^ SCs surrounded by ACTA2^+^ peritubular myoid cells (Fig. 2c, d). TFAP2C^+^ GCs were not present, suggesting the successful depletion of mPGCs (Fig. 2d). Tubules were surrounded by NR2F2^+^ stroma reminiscence of mouse fetal testis, but CD31^+^ endothelial cells and HSD3B1^+^ Leydig cells were not observed (Fig. 2d). Next, we sought to identify culture conditions that would allow the integration of hPGCLCs into rTestes while maintaining the testicular tissue integrity and maximizing the recovery of hPGCLCs at 14 days of ALI culture. Floating aggregates cultured using Knockout Serum Replacement (KSR)-based media formed larger aggregates with less surrounding cell debris and enhanced recovery of live cells compared with those cultured in fetal bovine serum (FBS)-based media (Fig. 2e, f). Addition of Y-27632, a potent inhibitor of Rho-associated, coiled-coil containing protein kinase (ROCK) during floating culture further enhanced formation of tighter and slightly larger aggregates and increased recovery of AG positive cells (Fig. 2e, f)^38^. Moreover, 14 days of ALI culture using KSR-but not FBS-based medium showed the formation of distinct tubules with integration of AG^+^TFAP2C^+^ hPGCLC-derived cells within tubules as revealed by IF analysis (Fig. 2e, g, h). IF analysis also confirmed that hPGCLCs maintained markers of PGCs (POU5F1, NANOG, SOX17 and TFAP2C) after 14 days of ALI culture (Fig. 2h). All GCs were uniformly labeled by AG, suggesting that they were all derived from hPGCLCs and not from endogenous mouse PGCs (Fig. 2g, h). Also, of note, Leydig cells and endothelial cells were not readily observed in aggregates, suggesting that these cell types might have different culture requirement for survival and/or proper differentiation (Fig. 2h). Overall, these findings confirmed that dissociated mouse testicular somatic cells and hPGCLCs can be self-assembled to form mini-fetal testicular tissues, which we named xenogeneic reconstituted testis (xrTestis).

### Extended culture of xrTestis

We next cultured xrTestes for a prolonged period to determine if hPGCLCs could be further differentiated into more advanced male GCs (Fig. 3a). IF analysis for xrTestis cultured for 42 and 77 days revealed persistence of tubular structures consisting of SOX9 SCs and AG^+^ hPGCLC-derived cells that remained predominantly localized within tubules (Supplementary Fig. 3a, b). While essentially all GCs in d42 xrTestes showed an early PGC phenotype (AG^+^/TFAP2C^+^/DDX4^-^/DAZL^-^), many of the AG^+^TFAP2C^+^ GCs at d77 xrTestes were strongly immunoreactive for the M marker, DAZL and somewhat more weakly for DDX4 and VT (Supplementary Fig. 3b, c). These cells exhibit slightly larger and more euchromatic nuclei with low-DAPI intensity and prominent nucleoli, characteristic of primate PGCs (Fig. 3b)^15^. Consistently, cells exhibited low levels of global DNA methylation as revealed by IF for 5-methylcytosine (5mC) (Supplementary Fig. 3d). These cells also retained pluripotency-associated markers, such as POU5F1 and NANOG (Fig. 3c, d), suggesting that most hPGCLC-derived cells had progressed towards the M stage but had not yet differentiated into the T1 stage. In keeping with these findings, flow cytometry analysis at d81 showed the emergence of AG^+^VT^+^ and AG^-^VT^+^ GCs within the dissociated xrTestes, representing 7.9% or 1.7% of total cells (62% or 13% of all human GCs), respectively (Fig. 3e).

**Fig. 3.**
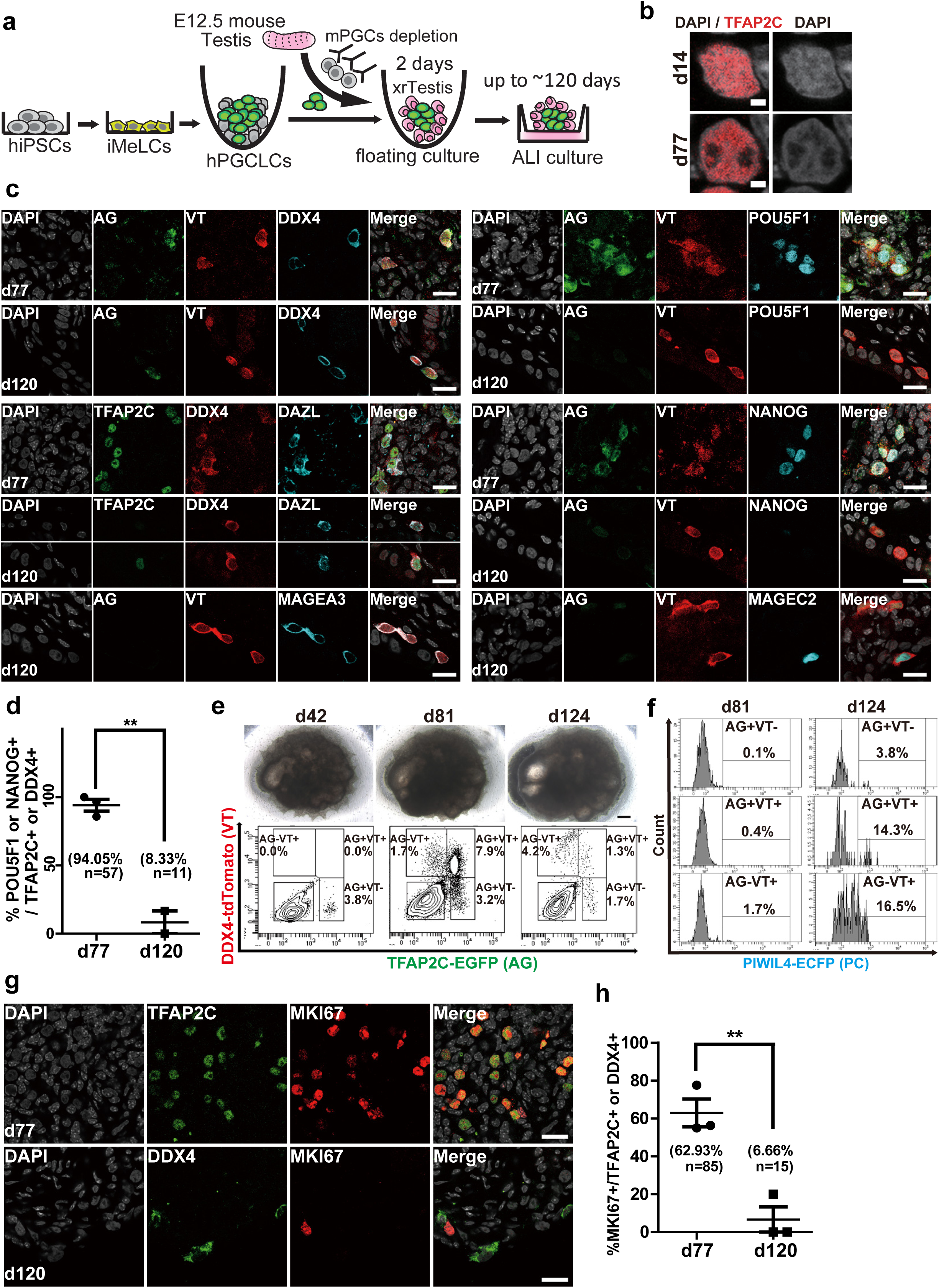
Establishment of xenogeneic reconstituted testis (xrTestis) and generation of human T1LCs. **a** Scheme for T1LC induction by xrTestis culture. ALI, air-liquid interphase. **b** IF images of hPGCLC-derived cells in day (d) 14 and d77 xrTestes for DAPI (white) and TFAP2C (red) with their merges. Bar, 2 µm. **c** IF images of d77 or d120 xrTestes for their expression of indicated key proteins (cyan: DAZL, POU5F1, NANOG, MAGEA3 and MAGEC2) in hPGCLC-derived cells [green: TFAP2C, TFAP2C-EGFP (AG) or red: DDX4, DDX4-2A-tdTomato, (VT)] and DAPI (white). Merged images are shown on the right. Bars, 20 µm. **d** A dot plot showing the proportion of POU5F1^+^ or NANOG^+^ cells in hPGCLC-derived cells (TFAP2C^+^ or DDX4^+^) in d77 and d120 xrTestes as assessed by IF analysis on frozen sections. Each dot represents the proportion in one section and the averages are shown. n, the number of positive cells. The statistical significance of the differences between d77 and d120 are evaluated by Fisher’s exact test. **P < 0.001. **e** Bright field (BF) images (top) and FACS analysis for AGVT expression (bottom) of xrTestes at d42, d81 and d124. Bar, 200 µm. **f** FACS histogram of d81 and d124 xrTestes for PIWIL4-ECFP (PC) expression in the respective fraction of hPGCLC-derived cells. % of PC+ cells in the respective fraction are shown. **g** IF images of hPGCLC-derived cells (TFAP2C or DDX4, green) in d77 or d120 xrTestes for MKI67 (red). Merges with DAPI (white) are shown on the right. **h** A dot plot showing the proportion of MKI67^+^ cells in hPGCLC-derived cells (TFAP2C^+^ or DDX4^+^) in d77 and d120 xrTestes. Each dot represents the proportion in one section and the averages are shown. n, the number of positive cells. The statistical significance of the differences between d77 and d120 are evaluated by Fisher’s exact test. **P < 0.001. See also Supplementary Fig. 2 and 3.

We next extended our ALI culture up to ∼d120: IF analysis on sections of xrTestes at this time revealed scattered DDX4^+^/DAZL^+^/VT^+^ cells with markedly reduced reactivity for AG, TFAP2C, POU5F1 or NANOG, suggesting that these cells had further differentiated beyond the M stage (Fig. 3c, d). We also noted the strong expression of MAGEA3 and MAGEC2, markers for T1 as defined above or elsewhere (Fig. 3c)^39^. Flow cytometric analysis confirmed that the majority of GCs had progressed to the AG^-^VT^+^ fraction (4.2% of total cells, 58% of all human GCs) (Fig. 3e). Moreover, we noted that some of the AG^+^VT^+^ (14.3%) and AG^-^VT^+^ cells (16.5%) expressed PC, a discriminating marker of T1 (Fig. 3f).

Finally, IF analysis showed the marked reduction of MKI67^+^ GCs in d120 xrTestes compared with those in d77 xrTestes, which is consistent with progression into the mitotically arrested T1 state (Fig. 3g, h). Together, these findings support our conclusion that hPGCLCs became T1-like cells (T1LCs).

### Lineage trajectory leading to T1LCs

To capture the precise continuum of male germ lineage progression of the late GCs from xrTestes without bias introduced by a priori markers, we isolated GCs (AG^+^ or VT^+^) from xrTestes at days 81 and 124 and evaluated their transcriptomes by scRNA-seq. Precursor cell types, hiPSCs, iMeLCs and hPGCLCs were also examined individually by scRNA-seq. Between 344-1251 cells for each population were used after excluding results from low quality cells (Supplementary Fig. 4a). We detected ∼3000-7000 median genes/cell at a sequencing depth of between 68-273k mean reads/cell (Supplementary Fig. 4a). After computational aggregation of high-quality cells and presentation in the same tSNE space (Fig. 4a, Supplementary Fig. 4b, c, f), we found 4 main clusters (cluster 1-4), which were largely separated by sample type, as expected (Supplementary Fig. 4b, f, g). Cluster 4 (yellow) consisted of GCs from both d81 and d124 xrTestes (Supplementary Fig. 4b, f). Cluster annotation was further validated by the expression of the known markers for each cell type such as *SOX2* (marker for hiPSCs and iMeLCs), *EOMES* (maker for iMeLCs), *NANOS3* (marker for hPGCLCs and later stage GCs) (Supplementary Fig. 4b)^17, 18^.

**Fig. 4.**
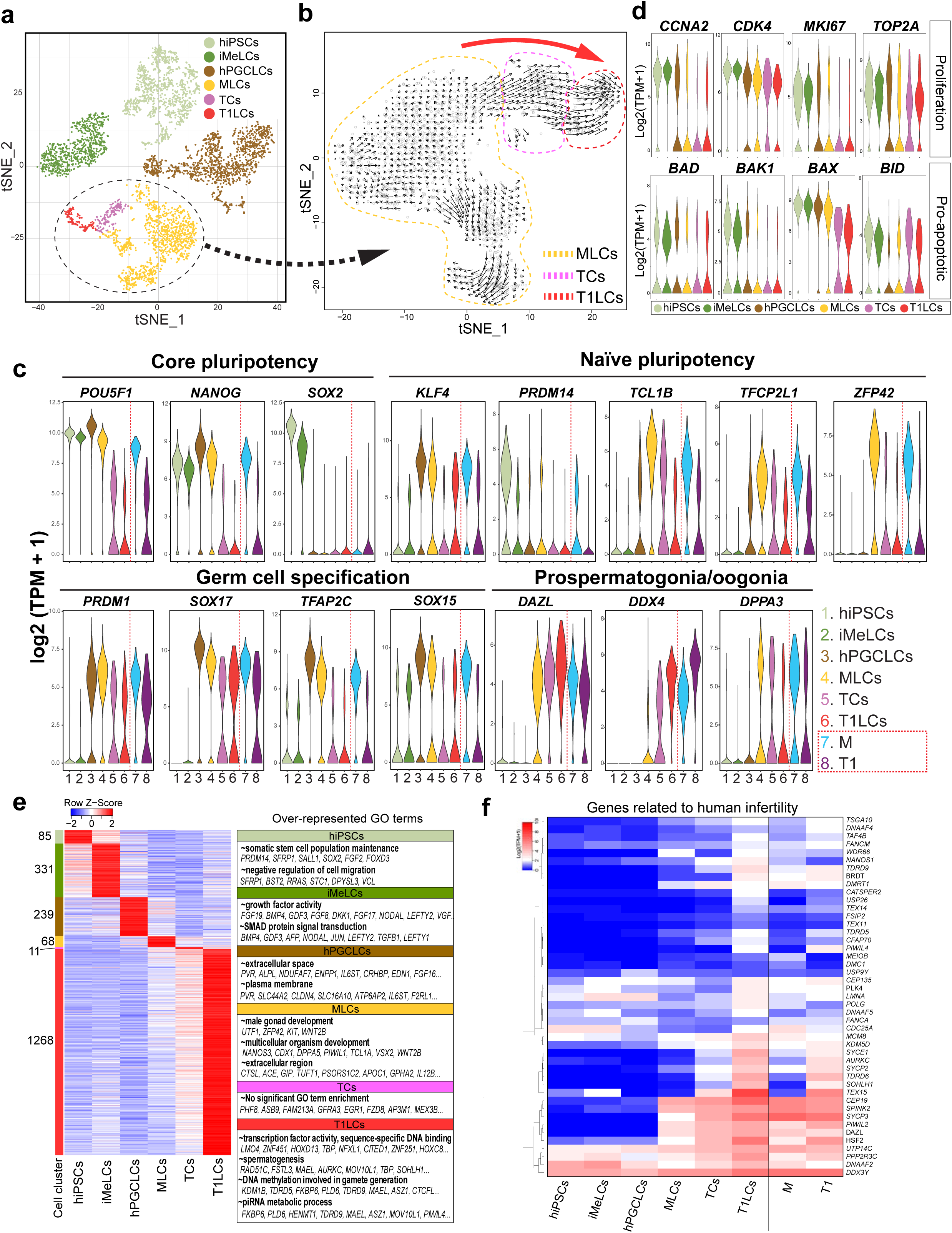
Single cell transcriptome profiling of *in vitro* derived human GCs. **a** A tSNE plot of all *in vitro* cells using computationally aggregated scRNA-seq data obtained from 6 different samples (hiPSCs; iMeLCs; hPGCLCs_1; hPGCLCs_2, d81 xrTestis, d124 xtTestis). These cells are colored based on clusters defined in Supplementary Fig. 4b and c: hiPSCs (1168 cells, pale green); iMeLCs (800 cells, green), hPGCLCs (1371 cells, brown), MLCs (1224, yellow), TCs (162 cells, purple) and T1LCs (150 cells, red). A cluster re-analyzed in (**b**) are outlined by dotted line. **b** Velocity analysis focused on a cluster outlined by dotted line in (**a**) projected on the tSNE space defined in Fig. S4E. Black arrows indicate the lineage trajectory estimated using nascent transcripts from scRNA-seq data in (**a**). Dotted lines enclose cells representing MLCs (yellow), TCs (purple) or T1LCs (red) as defined in Supplementary Fig. 4e. Large red arrow indicates overall direction of the lineage trajectory at the borders of cell clusters. **c** Violin plots showing the expression of representative markers in *in vitro* (left) or *in vivo* (right) cell clusters defined in Fig. 1a and 4a. **d** Violin plots showing the expression of representative proliferation markers (top) or pro- apoptotic markers (bottom) in the respective cell clusters. **e** Heatmap showing the expression pattern in respective cell clusters of DEGs obtained by multi-group comparison (FDR <0.01, fold change >2 compared to other clusters) and over-represented GO terms in these DEGs. Color bars at the left side of the heatmap indicate DEGs for respective cell clusters. The number of DEGs for each cell cluster are shown at the side of the color bar. **f** Heatmap of the expression of 45 genes, in which mutations are associated with “spermatogenic failure” or “male infertility” in humans. Genes are ordered by UHC (ward method). See also Supplementary Fig. 4-6 and Supplementary Table 2.

We noted that cells derived from xrTestes in cluster 4 expressed *DAZL, DND1, PIWIL2*, which was consistent with their prospermatogonial status (Supplementary Fig. 4b). Further analysis for cluster 4 in isolation revealed three distinct subclusters (Supplementary Fig. 4c). Markers for M defined previously *in vivo*, such as *KHDC3L* and *POU5F1* were uniquely expressed in cluster 4-1: therefore, we termed cells representing this cluster as “M-prospermatogonia-like cells (MLCs)”. Markers for T1 such as *MAGEC2* and *PIWIL4* were exclusively expressed in cluster 4-3, suggesting that this cluster represented T1LCs (Supplementary Fig. 4c). We noted a cluster located between T1LCs and MLCs, expressing unique markers, *ASB9* and *FZD8* (Supplementary Fig. 4c). We posited that these cells were transitional cells (TCs) because of their position in the tSNE plot and the intermediate levels of expression for both MLC (*POU5F1*, *NANOS3*) and T1LC markers (*TEX15*) (Supplementary Fig. 4c). Consistent with this interpretation, strong ASB9 expression was also seen in cells present at the border of M and T1 in a tSNE plot for *in vivo* human testicular GC samples evaluated previously (Supplementary Fig. 4d).

To understand the lineage trajectory among cells in cluster 4, we performed RNA velocity analysis after re-clustering (Fig. 4b and Supplementary Fig. 4e). This analysis confirmed that lineage progression from *POU5F1*^+^ MLCs to *PIWIL4*^+^ T1LCs occurred in xrTestis cultures (Fig. 4b). In line with this, the proportion of cells derived from d124 xrTestis increased from 10.2% in the MLCs cluster to 74.1% in the TC cluster and up to 99.3% in the T1LC cluster (Supplementary Fig. 4g). We also noted two biological replicates for hPGCLCs (hPGCLCs_1, hPGCLCs_2) were intermingled to form a single hPGCLC cluster in the tSNE plot (Supplementary Fig. 4f) and showed high concordance of averaged gene expression using whole genome data (r2=0.98) (Supplementary Fig. 4h) or pair-wise DEGs between hPGCLCs and MLCs (data not shown). This underscored the robustness of the present scRNA-seq platform.

To more fully understand the molecular mechanisms driving the differentiation of hT1LCs from hPGCLCs, we explored the gene expression dynamics of these *in vitro* derived GCs. The core pluripotency-associated genes, *POU5F1* and *NANOG*, along with GC specifier genes (*PRDM1, SOX17, TFAP2C, SOX15*) showed peak expression levels at the hPGCLC stage and declined thereafter (Fig. 4c). Interestingly, expression of some genes involved in naïve pluripotency such as *TCL1B, TFCP2L1, and ZFP42* peaked at the MLC stage. Consistent with our IF and flow cytometric data, prospermatogonial (or oogonial) markers, such as *DAZL, DDX4* and *DPPA3* emerged in MLCs and persisted in later GCs (Fig. 4c)^14, 27^. Moreover, in keeping with IF analysis (Fig. 3g, h), proliferation markers, such as *CCNA2*, *CDK4*, *MKI67* and *TOP2A* were significantly downregulated in T1LCs. Expression of proapoptotic marker genes were downregulated or unchanged during T1LC specification, suggesting that these cells were quiescent but not overtly apoptotic (Fig. 4d).

Our pairwise and multigroup DEG analyses showed the expression dynamics of genes involved in biologic processes that are unique to each cell type (Fig. 4e, Supplementary Fig. 5, 6, Supplementary Table 2). For example, iMeLCs exhibited higher abundance of mRNAs encoding gene products involved in early mesodermal specification, such as *EOMES, MIXL1, CER1* and genes related to SMAD signaling (e.g. *NODAL, LEFTY2*). During iMeLC-to-hPGCLC transition, hPGCLCs acquired expression of GC specifier genes along with genes enriched for GO terms related to inflammation and apoptosis (Supplementary Fig. 5 and Supplementary Table 2). Upon further differentiation into MLCs, some genes involved in “piRNA pathways” and “spermatogenesis” were turned on (Supplementary Fig. 5). We noted that T1LCs exhibited higher mRNA levels of the largest number of genes (1268 DEGs in multi-group comparison) and many of these DEGs were upregulated gradually during the transition from MLCs to T1LCs (Fig. 3e and Supplementary Fig. 6). In particular, we found genes involved in piRNA pathways and previously recognized markers for spermatogonia in DEGs for T1LCs (Fig. 4e, Supplementary Fig. 5, 6, Supplementary Table 3). Importantly, numerous transcription factors were also included in this list of DEGs, such as *LMO4, ZNF451, TBP, NFXL1, EGR4, SCX, EMX1,* many of which were previously uncharacterized in the context of male germline development and may play important roles in the specification of prospermatogonia from PGCs (Fig. 4e, Supplementary Fig. 5, 6, Supplementary Table 3). In keeping with this observation, these DEGs were enriched for those bearing GO terms, such as “transcription factor activity”, “spermatogenesis”, and “DNA methylation involved in gamete generation” (Fig. 4e). We also found that a significant fraction of genes known to be associated with human “spermatogenic failure” or “male infertility” were in T1LCs, suggesting the potential utility of this *in vitro* platform to dissect out the molecular mechanisms of human male infertility (Fig. 4f).

### T1LCs exhibit a transcriptome similar to that of T1 *in vivo*

With transcriptome data from both *in vivo* and *in vitro* samples in hand, we next set out to compare them side by side to precisely define the status of our *in vitro* derived GCs in the developmental coordinate. All three *in vivo* testicular samples were intermingled in a tSNE plot and segregated into clusters representing biologically meaningful cell types rather than batches (Supplementary Fig. 1c, g). Nonetheless, a scatter plot of the averaged gene expression for either whole GCs or T1 among three different samples revealed a modest genome-wide reduction of mRNA abundance in Hs26 and Hs27 (cryopreserved-thawed) compared with Hs31 (fresh) (Supplementary Fig. 1i), likely a technical consequence. Hs31 contained both M and T1 similar to aggregated sample (Supplementary Fig. 1h), and pairwise DEGs between M and T1 from Hs31 was similar to those obtained from the aggregation of all 3 samples (Supplementary Fig. 1j, k). Therefore, to compare gene expression in high precision with *in vitro* samples, which were all prepared fresh, we decided to use only Hs31for downstream comparative studies.

Unsupervised hierarchical clustering using averaged expression values for each cell type showed two large clusters, a pre-germ cell/pre-gonadal phase cluster (hiPSCs, iMeLCs, hPGCLCs) and a prospermatogonial (gonadal) phase cluster (M, MLCs, TCs, T1, T1LCs) (Fig. 5a). Within the latter cluster, M/MLCs and T1/T1LCs were each clustered together. Principal component analysis (PCA) focusing on prospermatogonial phase clusters revealed a gradual transition of cellular properties from M/MLCs to T1/T1LCs via TCs (Fig. 5b). Pairwise comparison of cell types along this lineage trajectory demonstrated high concordance; MLCs and T1LCs had the lowest number of DEGs and highest coefficient of determination (r^2^) when compared with M or T1, respectively (Fig. 5c). Together, these results highlighted the marked similarities of the progression of the fetal male germline developmental program *in vitro* and *in vivo*. Nonetheless, we also noted modest differences in the gene expression signatures between T1 and T1LCs, and in particular, noted that expression of some *HOXA* and *HOXB* family members was significantly higher in T1LCs (Fig. 5d and Supplementary Table 3). Reciprocally, *XIST*, a master regulator of X chromosome inactivation, expressed in human oogonia and prospermatogonia^40^, was expressed at significantly higher levels in *in vivo* T1 compared with T1LCs, as were 18 other genes (Fig. 5d). We next compared these findings with previously published datasets derived from human male fetal GCs which were categorized into migrating, mitotic and mitotic-arrest FGCs^14^. In these datasets, migrating FGCs were previously collected from Aorta-gonad-mesonephros region of human male embryos at the age of 4 weeks and therefore were categorized as PGCs in our study. The differentially-expressed genes that distinguished these three cell types separated our hPGCLCs, M/MLCs, and T1/T1LCs/TCs largely into “migrating”, “mitotic” and “mitotic-arrest” stages of human FGCs, respectively, in agreement with direct comparison of our *in vivo* and *in vitro* specimen (Fig. 5e, Supplementary Table 4). Finally, plotting of all the clusters in our dataset with previously published neonatal human spermatogonia within the same PCA coordinate showed that T1LCs are slightly closer than T1 to neonatal spermatogonia (Supplementary Fig. 4i, j), which might partly explain the differences we observed between T1 and T1LCs (Fig. 5d).

**Fig. 5.**
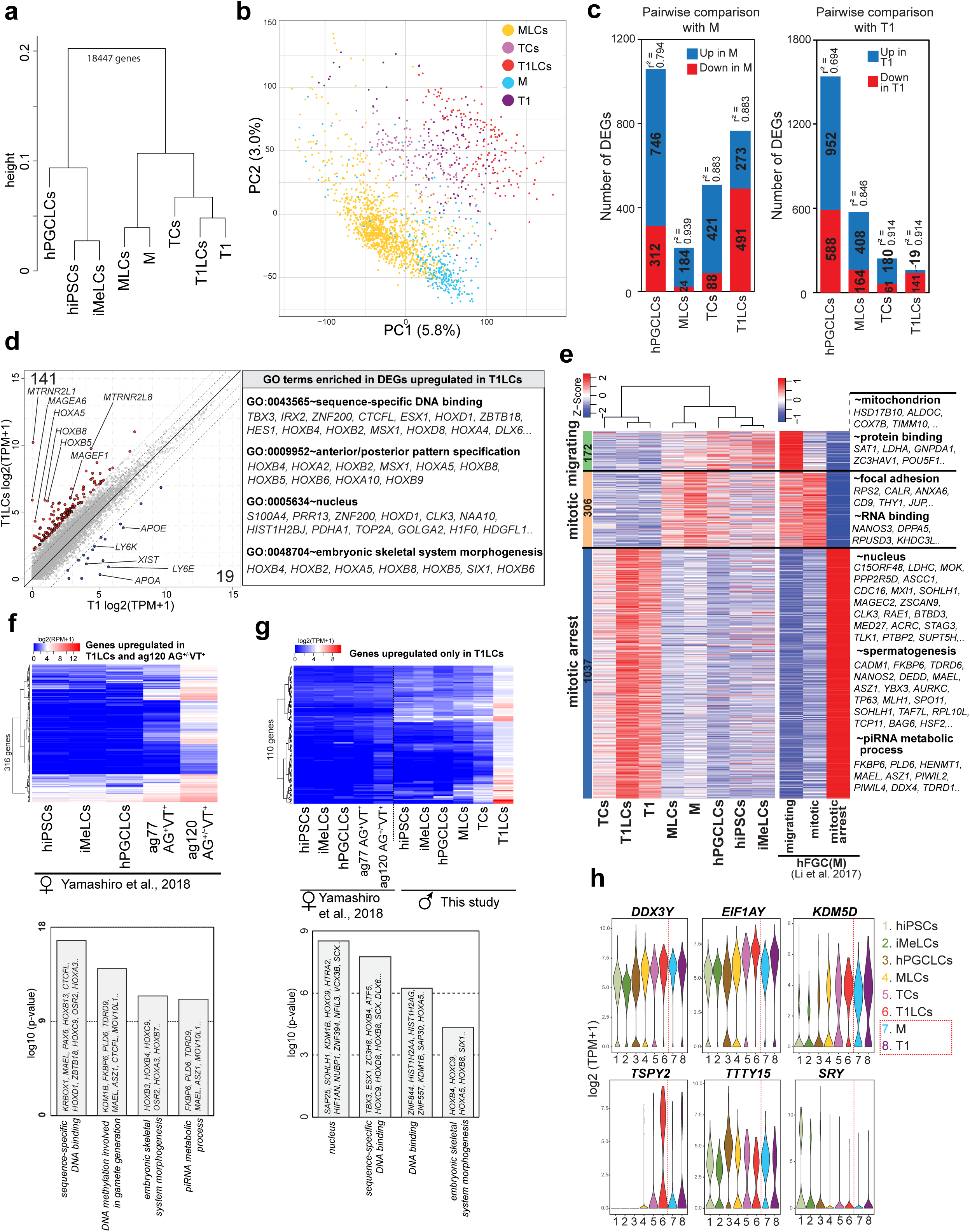
Comparison of lineage projection *in vitro* with that of *in vivo*. **a** UHC of the averaged transcriptomes of *in vivo* (M, T1) and *in vitro* cell clusters (hiPSCs, iMeLCs, hPGCLCs, MLCs, TCs, T1LCs) defined in Fig. 1a and 4a. **b** PCA for prospermatogonial stage GCs (M, MLCs, TCs, T1, and T1LCs). Color codes for cell clusters are indicated. **c** Number of DEGs and coefficient of determination (r^2^) by pairwise comparisons between *in vivo* [M (left) or T1(right)] and *in vitro* clusters (hPGCLCs, MLCs, TCs and T1LCs). **d** Scatter plot comparison of the averaged values of gene expression between T1 and T1LCs. blue, genes higher in T1; red, genes higher in T1LCs (more than 4-fold differences [flanking diagonal lines], mean log2(TPM+1) >2, FDR <0.01). Key genes are annotated and the number of DEGs are indicated. Representative genes and their GO enrichments for genes higher in T1LCs are shown on the right. **e** Heatmaps of the expression of markers for “migrating (172 genes)”, “mitotic (306 genes)” and “mitotic arrest (1037 genes)” human male (f)FGCs (defined by Li et al., 2017) in respective cell clusters defined in this study (left) and indicated hFGC types defined by Li et al., 2017 (right). Using these markers, hierarchical clustering is performed for cell clusters defined in this study (left). GO terms for the respective markers are shown on the right. **f** Heatmap of the averaged expression values of 316 genes in indicated cell types defined by Yamashiro et al., 2018 (top). Among markers for T1LCs defined in Fig. 4e, 1177 genes were also annotated in the dataset by Yamashiro et al., 2018. Among them, 316 genes upregulated in ag120 AG ^+/-^VT^+^ oogonia-like cells compared with all other 4 cell types (more than 2-fold difference) are used (top). GO analysis for these 316 genes are also shown (bottom). **g** Heatmap of the averaged expression in the indicated cell types for 110 genes highly expressed in T1LCs but have weak/no expression in ag120 AG^+/-^VT^+^ oogonia-like cells. To enable comparison between two different scRNA-seq platform, RPM values of the data from Yamashiro et al. were adjusted using the polynomial regression curve (y= 0.0908×2-0.0454x+0.1243, least square method) defined using scatter plot comparison between hiPSCs samples from this study and Yamashiro et al. Among 1177 DEGs for T1LCs (Fig. 4e), 110 genes showing high levels of expression in T1LCs [mean log2(TPM+1) > 4] and low levels of expression in oogonia-like cells [adjusted log2(RPM+1) < 2] are shown. GO analysis for these 110 genes are also shown (bottom) **h** Violin plot showing the expression levels of non-pseudoautosomal Y chromosome genes in indicated cell clusters. These genes are identified by multi-group DEG analysis in Fig. 4e (without cut-off by fold-change >2). See also Supplementary Fig. 4-6, Supplementary Table 3-5.

### Comparison of T1LCs with oogonia-like cells induced from hiPSCs

We next compared the transcriptome profiles of T1LCs with ag120 AG^+/-^VT^+^ oogonia-like cells induced from hiPSCs using a published database^18^. Among the 1177 DEGs unique to T1LCs and commonly annotated in both platforms, 316 were also similarly higher in oogonia-like cells (Fig. 5f and Supplementary Table 5), including many genes involved in piRNA biogenesis and *de novo* DNA methylation. We also identified 110 genes that were highly expressed in T1LCs but had weak/no expression in oogonia-like cells (Fig. 5g and Supplementary Table 5). These genes included many transcription factors, such as *HOXB4, HOXC9, TBX3, ESX1, SCX, SIX1*, and accordingly were enriched with those bearing GO terms such as “nucleus” and “sequence-specific DNA binding.” Of note, our T1LCs were induced from hiPSCs bearing an XY karyotype. Accordingly, we found that 6 non-pseudo-autosomal, Y chromosomal genes were differentially expressed among our *in vitro* dataset (Fig. 5h). Among these genes, we noted that 4 genes, *DDX3Y, EIF1AY, KDM5D, TSPY2* showed a significant upregulation along the lineage trajectory, with a similar trend also being observed between M and T1. These findings highlighted the commonality and differences between male and female fetal GC development.

### Expression dynamics of transposable elements during human male germline development

Activation and the subsequent repression of transposable elements (TEs) is a hallmark feature of GC development, which ensure genome integrity of GCs while permitting co-evolution between TEs and their hosts^19, 41–43^. However, the precise expression dynamics of TEs in human male GC development is not well characterized. Taking advantage of our *in vitro* platform that covers at least the first ∼20 weeks of human male GC development, we went on to comprehensively map the expression patterns of various TE categories. Dimension reduction analysis of scRNA-seq data based on TE expression profiles demonstrated that each differentiation stage exhibits unique TE expression dynamics (Fig. 6a). Notably, M/MLCs and T1/T1LCs were clustered together in a UMAP plot and a hierarchical clustering using variably expressed TEs, further underscoring the similarities (Fig. 6a, b). Analysis of the expression of total TE-derived transcripts showed global upregulation of TEs over time, both *in vivo* and *in vitro* with the highest expression levels observed in T1 and T1LCs (Fig. 6c). Consistently, most TE subtypes belonging to LINE1, SINE, DNA transposon, and SVA showed gradual derepression, presumably due to a genome-wide DNA demethylation and other epigenetic reprogramming events that accompany fetal GC development (Fig. 6d, e)^27, 40, 42^. In contrast to the above TEs, LTR-type retrotransposons (i.e., endogenous retroviruses; ERVs) tended to show more stage-specific expression pattern (Fig. 6d-f). For example, among ERV1 families, HERV-H and LTR7 expression peaked at hiPSCs, with a subsequent gradual decline in expression levels (Fig. 6f), consistent with their roles in regulating the pluripotency gene network^44–46^. Another group of ERV1 families, the LTR12 families (LTR12C, LTR12D, LTR12_) were significantly upregulated in T1 or T1LCs (Fig. 6f). On the other hand, some members of the ERVK family, such as HERVIP10FH were more highly expressed in hPGCLCs (Fig. 6f).

**Fig. 6.**
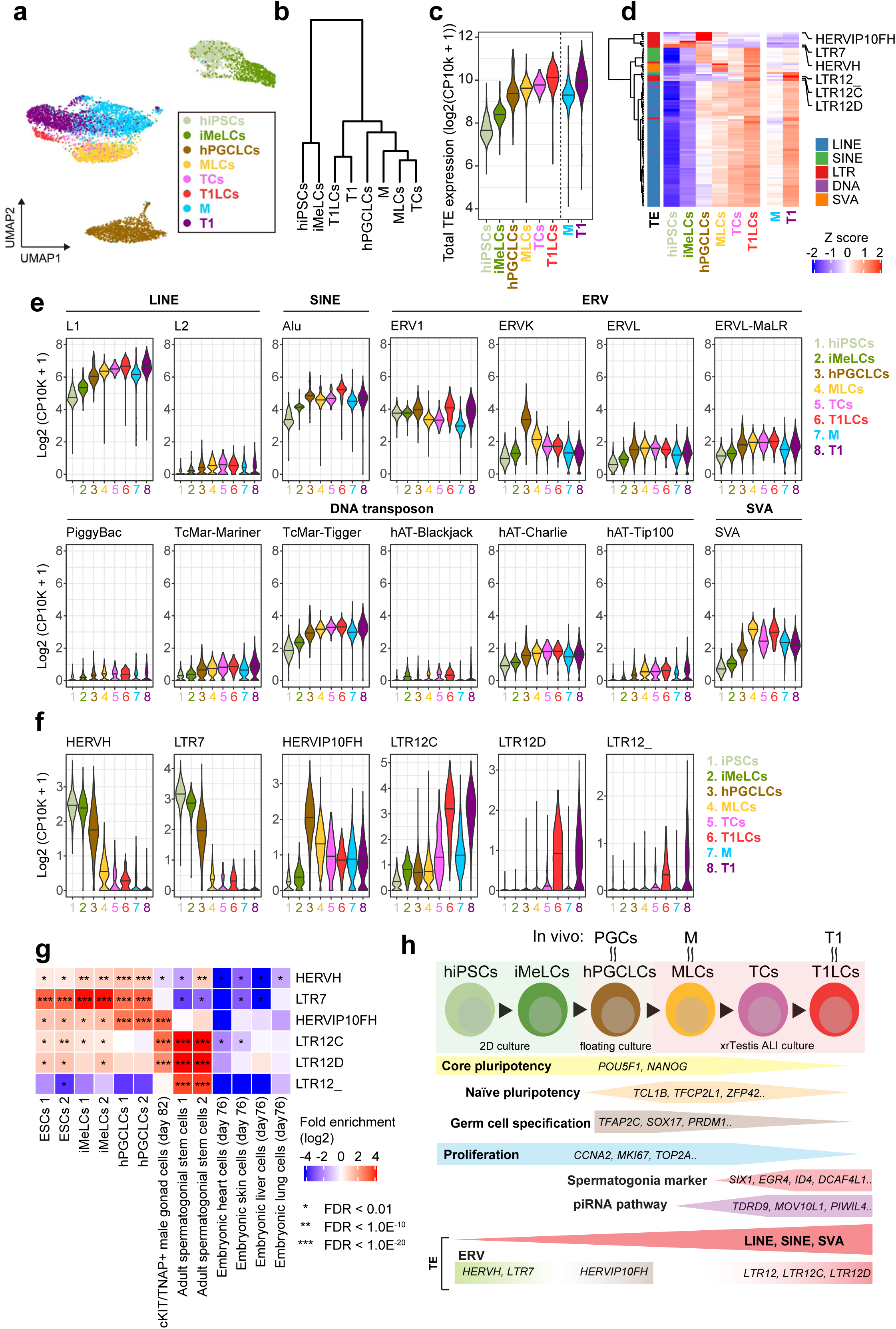
Dynamic regulation of TE expression during human male germline development. **a** Dimension reduction analysis of scRNA-Seq data based on TE expression profile. Most variably expressed 100 TE subfamilies are used in the analysis. **b** Hierarchical clustering (ward method using Euclidian distances) of the respective cell clusters (averaged expression values) using most variably expressed 200 TEs. **c** Dynamics of the total expression level of TEs. **d** Heatmap showing the expression dynamics of respective TE subfamilies. Most variably expressed 200 TEs are shown. Classification of TEs are shown in the left of heatmap. **e** Expression dynamics of respective TE families. **f** Expression dynamics of ERV subfamilies. ERVs particularly exhibiting unique expression patterns are shown. **g** Enrichment of ATAC-Seq signals on specific ERV subfamilies. The fold enrichment of the overlaps between ATAC-Seq peaks and a specific ERV subfamily was calculated on the random expectation. Publicly available ATAC-Seq data (Chen et al., 2018 and Guo et al., 2017) were used. **h** A model for T1LC induction from hiPSCs using xrTestis culture.

Stage-specific expression dynamics of ERVs implicates them in regulation of the expression of genes involved in development. In fact, the analysis of publicly available ATAC-seq data revealed the enrichment of HERVH/LTR7 and HERVIP10FH in the ATAC-seq peaks obtained from ESCs/iMeLCs or hPGCLCs, respectively (Fig. 6g), coinciding with the expression peaks in respective cell types (Fig. 6f). Moreover, LTR12 subfamilies were enriched in the ATAC-seq peaks from human male gonadal GCs and adult spermatogonia suggesting that these TEs might involve in gene regulation in human prospermatogonia/spermatogonia (Fig. 6g).

In summary, our study demonstrating that ERVs are regulated in a dynamic and stage-specific manner raise the intriguing possibility that ERVs may be directly involved in the transcriptional activation of genes driving human prospermatogonial specification and suggest that identification of sequences downstream of these regulatory elements may identify genes that play a causative role in this process (Fig. 6h).

## DISCUSSION

A significant number of male infertility are due to SCOS resulting from the permanent loss of the male germline because of errors during fetal or postnatal development or iatrogenic insults such as chemotherapy and radiation treatments for cancer^47^. The diagnosis and treatment of SCO is currently limited by our lack of understanding of the mechanistic basis driving the developmental progression of the human male germline to ultimately enable spermatogenesis. Since adult spermatogenesis is carried out by SSCs with perpetual self-renewal capability, treatment of SCOS could be potentially executed by establishing human spermatogonial cultures for gamete derivation either *in vitro* or *in vivo* following transplantation. In mice, cells possessing SSC activities can be propagated perpetually, allowing for their characterization, genetic manipulation, and transplant into infertile recipients to replenish spermatogenesis^48, 49^. However, despite numerous attempts by many laboratories, sustained human spermatogonial cultures have yet to be convincingly established^49^. Moreover, defective gametogenesis in many infertile men occurs at a pre-meiotic stage, precluding the use of SSC-dependent reconstitution systems. However, this limitation could be overcome if male GCs could be derived from hiPSCs after correction of causative mutations *in vitro*. Derivation of functional male GCs from iPSCs was recently accomplished by several groups in mice^37, 50^. In this study, we provide the first evidence that hiPSCs can be used to reconstitute fetal human male germline development *in vitro* through stepwise and faithful recapitulation of the normal developmental processes (Fig. 6h).

We found that in xrTestes, ∼75% hPGCLC-derived cells differentiated into VT^+^ MLCs or TCs by ∼d80 of ALI culture (Fig. 3e). Although we did not further refine the exact timing at which the hPGCLC-to-MLC transition occurs, the developmental kinetics of GCs in xrTestis appears to be somewhat faster than that of xrOvary, in which only a small fraction (∼15%) showed VT expression by ∼d80 of ALI culture^18^. Moreover, the AG^-^VT^+^ fraction, which seems to be enriched for GCs at advanced stages (TCs, T1LCs or RA-responsive oogonia-like cells) contained ∼13% of GCs in xrTestis at ∼d80. In contrast, the AG^-^VT^+^ fraction only emerges after ∼d100 in xrOvaries and at lower frequency (∼3%), suggesting that xrTestes might provide a better niche environment than xrOvaries for sex-specific GC progression into advanced stages. Future studies directly comparing the two culture platforms using the same hiPSC lines of both XX and XY karyotypes may be warranted to determine if this finding is universal.

In extended cultures of xrTestes, we noted the dissolution of the majority of tubular structures after ∼day 80, and as a result, GCs were randomly localized in the stroma with apparent loss of contact with SCs by d120. However, the fate of T1LCs appeared to be maintained until at least d120, even in the absence of tubular structure. The absence of a tubular structure after d120 will likely pose significant challenges for further differentiation of T1LCs into more advance male GC types. In particular, it may be difficult to derive neonatal or adult spermatogonia which require continuous interaction with the appropriate niche environment, particularly that provided by SCs^20–22^. The underlying reasons for tubular dissolution are still unclear but may be due to the different nutritional requirements for the long-term cultured SCs, which may more closely reflect their postnatal counterparts late in our culture period. Lack of endothelial cells in xrTestes evident as early as d14 (Fig. 2) might also contribute to this dissolution given the well-established role of the vasculature in testicular cord formation and proper structural testis development^51, 52^. Supplementing xrTestes with cultured testicular endothelial cells, or growth factors that sustain their survival might help vascularize xrTestes. In order to establish more robust, sustainable, and complete reconstitution of prospermatogonial development, future studies may also seek to replace the xrTestis constituents with more physiological niches, such as those derived from human fetal testis or their mimetics induced from hiPSCs.

Despite some limitations, our xrTestis culture approach is the first method that reliably produces T1 stage human fetal prospermatogonia from hiPSCs. Surprisingly, we found that T1LCs share a number of key genes previously defined to mark mouse or human adult spermatogonia, such as *SIX1, DCAF4L1, PLPPR3, EGR4*, or *ID4* among others (Fig. 5, Supplementary Fig. 5, 6, Supplementary Table 2)^20, 21, 35, 53^. Moreover, transcriptome comparison of T1LCs and T1 with neonatal prospermatogonia from a previous report suggested that T1LCs appear to be closer to neonatal prospermatogonia than are T1 (Supplementary Fig. 4). Due to the paucity of data currently available regarding perinatal human male germline development, the exact *in vivo* stage to which T1LCs correspond remains unclear. Recent studies suggested that SSC activity, the competency to produce self-sustaining spermatogenesis upon transplantation has already been established, at least in part in fetal prospermatogonia in mice^54^. Thus, it is possible that T1LCs might also be close to foundational SSCs which upon transplantation would be capable of regenerating complete spermatogenesis. Experimental validation of this possibility would require a relevant model, such as non-human primates, in which the full regenerative potential of T1LCs could be assessed^55, 56^.

T1LCs derived from our xrTestis culture exhibited a transcriptome signature in which many genes linked with human male infertility were specifically expressed, opening the door for using the xrTestis platform to dissect the cellular and molecular mechanisms of human male infertility (Fig. 4, 5). For example, among DEGs we identified, *DDX3Y* has been suggested to play a critical role in fetal male GC development and the mutation appears to be responsible for one of the common causes of SCOS caused by Y-chromosome microdeletions^57, 58^. T1LCs also specifically expressed genes involved in piRNA biogenesis (Fig. 4, 5, Supplementary Fig. 5, 6), suggesting that xrTestis platform may also prove to be useful for addressing the molecular mechanisms of piRNA production and consequent regulation of various TEs, fossilized viral descendants, that can have detrimental effects on genome integrity if not properly controlled during spermatogenesis^59–62^. Interestingly, recent studies highlight the key role of LTRs as cis-regulatory elements in the transcriptional regulation of genes involved in early embryogenesis, suggesting that host-viral coevolution exploited TEs that are typically upregulated during early embryo or germline development due to the epigenetically permissive genomic status^63–66^. In line with this notion, we showed that LTR12 subfamilies were uniquely expressed among T1LCs and T1s and are positionally enriched in ATAC-seq peaks identified in human prospermatogonia or spermatogonia, suggesting that these LTRs may serve as cis-regulatory elements for germline development (Fig. 6). Future studies perturbing expression of LTRs using CRISPRa or CRISPRi^64^ on xrTestes might permit functional interrogation of LTRs in human GC programming.

In sum, we provide the first demonstration that human fetal male GC development can be reconstituted from hiPSCs (Fig. 6g). As such, xrTestes will be useful resource for understanding various aspect of human male gametogenesis and a step forward for future production of human sperm in dish.

## METHODS

### Mice

Animal procedures were conducted in compliance with Institutional Animal Care and Use Committee of the University of Pennsylvania. Timed pregnant ICR female mice were purchased from Charles River (Wilmington, MD).

### Collection of human fetal samples

Fetal testis samples at 17-18 weeks of gestation were obtained from three donors undergoing elective abortion at the hospital of the university of Pennsylvania. All experimental procedures were approved by IRB (protocol#832470) and Human Stem Cell Research Advisory Committee at the University of Pennsylvania. Informed consent was obtained from all the human subjects.

### Culture of hiPSCs

The experiments on the induction of human primordial germ cell-like cells (hPGCLCs) from human induced pluripotent stem cells (hiPSCs) were approved by Institutional Review Board of University of Pennsylvania. hiPSCs were culture on a plate coated with Recombinant laminin-511 E8 (iMatrix-511 Silk, Nacalai USA) and were maintained under a feeder-free condition in the StemFit® Basic04 medium (Ajinomoto) containing basic FGF (Peprotech) at 37 °C under an atmosphere of 5% CO2 in air. For the passage or the induction of differentiation, the cells were treated with a 1 to 1 mixture of TrypLE Select (Life Technologies) and 0.5 mM EDTA/PBS for 14 min at 37 °C to enable their dissociation into single cells. 10 μM ROCK inhibitor (Y-27632; Tocris) was added in culture medium for 1 day after passaging hiPSCs.

### Human testis sample preparation

The sex of fetuses was determined by sex-specific PCR on genomic DNA isolated from mesonephros and attached fibroconnective tissues using primers for the ZFX/ZFY loci (Table S6)^67^. Fetal testes were dissected out in RPMI-1640 (Roche) and ∼one fourth of the tissues were fixed in 10% formalin overnight at room temperature before processing for histologic analyses.

The remaining testicular tissues were dissociated into single cells by two-step enzymatic digestion protocol^68^. Briefly, fetal gonads were cut into small pieces (∼1mm^2^), then digested with collagenase type IV for 8 min at 37 °C. Tissues were centrifuged at 200 g for 5 min and washed with Hank’s Balanced Salt Solution (Thermo Fisher Scientific). Tissues were then digested with 0.25% trypsin/EDTA and DNase I for 8 min at 37 °C. After the digestion was quenched by adding 10% of fetal bovine serum, cells were strained through nylon cell strainer (70μm in pore size). The cells were then pelleted by centrifugation at 200 g for 5 min, which were resuspended in the MACS buffer (PBS containing 0.5% BSA and 2 mM EDTA). Dead cells were subsequently removed by Dead cell removal microbeads (Miltenyi Biotec). Flow-through cells were pelleted by centrifugation at 600 g for 15 min and resuspended in PBS containing 0.1% BSA before loaded in 10x Genomics Chromium Controller.

### Generation of AGVTPC-Knock-in reporter lines

The donor vector and the TALEN constructs for generating the *DDX4/hVH-p2A-tdTomato* (VT) allele were described previously ^18^. Briefly, homology arms of *DDX4* (left arm:1471 bp; right arm: 1291 bp) were amplified by PCR from the genomic DNA of *TFAP2C-2A-EGFP* (AG) male hiPSCs (585B1 1-7 kind gift from Dr. Mitinori Saitou, Kyoto University) and were sub-cloned into the pCR2.1 vector using the TOPO TA cloning kit (Life Technologies). The *p2A-tdTomato* fragment with the PGK-Neo cassette flanked by *LoxP* sites was also amplified by PCR and inserted in-frame at the 3’-end of the *DDX4* coding region of the subcloned vector. TALEN constructs targeting *DDX4* were generated using a Golden Gate TALEN and TAL Effector kit 2.0 (Addgene, #1000000024)^69^. TALEN’s RVD sequences were as follows: DDX4-left (5-prime), HD HD HD NI NI NG HD HD NI NN NG NI NN NI NG NN NI NG NN; DDX4-right (3-prime), NN NI NI NN NN NI NG NN NG NG NG NG NN NN HD NG NG^18^.

To construct the donor vector for generating *PIWIL4-p2A-ECFP* (PC) allele, the homology arms of *PIWIL4* (left arm: 1,497bp; right arm: 1,000 bp) was amplified by PCR and was sub-cloned into pCR2.1 vector using TOPO TA cloning kit. The *p2A-ECFP* fragment with the *PGK-Puro* cassette flanked by *loxP* sites was synthesized by GeneUniversal (Newark, DE), and inserted in-frame at the 3’-ends of *PIWIL4* coding sequence of the sub-cloned vector using GeneArt Seamless Cloning & Assembly Kit (Life Technologies). The stop codon was removed to allow the expression of the in-frame p2A-ECFP protein. *MC1-DT-A-polyA* cassette was synthesized and subsequently inserted into the downstream region of the right homology arm of the PC donor vector using the restriction enzymes NotI/XhoI.

The single guide RNAs (sgRNAs) pair targeting the sequence close to the stop codon of *PIWIL4* were designed by Molecular Biology CRISPR design tool (Benchling). Two sgRNAs (GAACTACTGGCATCACTAGA and TCACAGGTAGAAGAGATGGT) were selected and cloned into pX335-U6-Chimeric BB-CBh-hSpCas9n (D10A) SpCas9n expressing vector, respectively to generate sgRNAs/Cas9n vector (Addgene, #42335)^70^. The recombination activity of the designed sgRNAs/Cas9n was validated by the single-strand annealing (SSA) assay.

The donor vectors (5 µg) and TALEN and sgRNAs/nCas9 vectors (2.5 µg each) for DDX4 and PIWIL4 were introduced into one million AG hiPSCs (585B1 1-7, male) by electroporation using the NEPA21 type II (Nepagene). Single colonies were picked up after selection with puromycin and neomycin and a subsequent transfection with a plasmid expressing Cre recombinase to remove *PGK-Puro* and *PGK-Neo* cassettes. The successful targetings, the random integrations and Cre recombinations were assessed by PCR on the extracted genomic DNA from each colony using the primer pairs listed in Table.S6.

### Karyotyping and G-band analyses

hiPSCs were incubated with the culture medium containing 100ng/ml of karyoMAX colcemid (Gibco) for 8 hr. After dissociation by TrypLE select, cells were treated with pre-warmed Buffered Hypotonic solution and incubated for 30 min at 37 °C. Cells were then fixed with Carnoy’s solution (mixture of methanol and acetic acid at 3:1 ratio) and dropped onto glass slides to prepare chromosomal spread. Karyotypes were first screened by counting the numbers of chromosomes identified by DAPI staining. Cell lines bearing 46 chromosomes were further analyzed by G-banding method performed by Cell Line Genetics (Madison, WI).

### Induction of hPGCLCs and the generation of xrTestes

hPGCLCs were induced from hiPSCs via iMeLCs as described previously^17^. For the induction of iMeLCs, hiPSCs were plated at a density of 4-5×10^4^ cells/cm^2^ onto a human fibronectin (Millipore)-coated 12 well plate in GK15 medium (GMEM [Life Technologies] with 15% KSR, 0.1 mM NEAA, 2mM L-glutamine, 1mM sodium pyruvate and 0.1mM 2-mercaptoethanol) containing 50 ng/ml of ACTA (R&D Systems), 3 μM CHIR99021 (Tocris Bioscience) and 10 μM of a ROCK inhibitor. After 31-38 hrs, iMeLCs were harvested and dissociated into single cells with TrypLE Select. Cells were then plated into a well of a low-cell-binding V-bottom 96-well plate (Thermo Fisher Scientific) at 3500 cells per well in GK15 medium supplemented with 200 ng/ml BMP4 (R&D Systems, 314-BP-010), 100 ng/ml SCF (R&D Systems, 255-SC-010), 50ng/mL EGF (R&D Systems, 236-EG), 1,000 U/ml LIF (Millipore, #LIF1005) and 10 μM of Y-27632 to be induced into hPGCLCs.

Xenogeneic reconstituted testes (xrTestes) were generated by aggregating FACS-sorted d5 hPGCLCs with fetal testicular somatic cells of E12.5 embryos according to a procedure described previously^37^. The d5 floating aggregates containing hPGCLCs, which were induced from AGVTPC hiPSCs, were dissociated into single cells with 0.1% Trypsin/EDTA treatment for 15 min at 37 °C. After quenching the reaction by adding an equal volume of FBS, cells were resuspended in FACS buffer (0.1% BSA in PBS) and were strained by a cell strainer (Thermo Fisher Scientific) to remove cell clumps. Then, AG^+^ cells were collected by FACS. To isolate fetal testicular somatic cells, E12.5 embryos were isolated from timed pregnant ICR females and collected in chilled DMEM (Gibco) containing 10% FBS (Gibco) and 100 U/ml penicillin/streptomycin (Gibco). Fetal male testes were identified by their appearance, and the mesonephros were removed by tungsten needles. Isolated E12.5 testes were washed with PBS and incubated with dissociation buffer containing 1mg/ml Hyaluronidase Type IV (Sigma), 5U Dispase (Corning), and 5U DNase (Qiagen) in wash buffer (100 U/ml penicillin/streptomycin and 0.1% BSA in DMEM/F12) for 15 min with periodical pipetting at 37 °C. After washing with PBS, testes were dissociated into single cells by 0.05% Trypsin-EDTA in PBS for 10 min at 37 °C, followed by quenching with FBS. Cell suspension was strained through a 70 µm nylon cell strainer, centrifuged at 1,200 rpm for 5 min, and the supernatant was discarded. The cell pellet was resuspended with MACS buffer (PBS containing 0.5% BSA and 2 mM EDTA), and then was incubated with anti-SSEA1 antibody MicroBeads (Miltenyi Biotec) for 20 min on ice. The cell suspension was washed with MACS buffer and centrifuged at 1,200 rpm for 5 min, and the supernatant was removed. The cell pellet was resuspended in MACS buffer and then applied to an MS column (Miltenyi Biotec) according to manufacture protocol. The flow-through cells were centrifuged at 1,200 rpm for 5 min, resuspended with Cell banker Type I (Amsbio, ZNQ CB011) and cryopreserved in liquid nitrogen until use.

To generate floating aggregates, hPGCLCs (5,000 cells per xrTestis) and thawed fetal testicular somatic cells (60,000 cells per xrTestis) were mixed and plated in a Lipidure-coated U-bottom 96-well plate (Thermo Fisher Scientific, 174925) in Minimum Essential Medium alpha (α-MEM) (Invitrogen) containing 10% KSR (Gibco), 55 µM 2-mercaptoethanol (Gibco), 100 U/ml penicillin/streptomycin (Gibco) and 10 µM Y-27632. After 2days of floating culture, floating aggregates were transferred onto Transwell-COL membrane inserts (Corning, 3496) soaked in α -MEM containing 10% KSR, 55 µM 2-mercaptoethanol, 100 U/ml penicillin/streptomycin using a glass capillary. xrTestes were cultured at 37 °C under an atmosphere of 5% CO2 in air. Half the volume of medium was changed every three days.

### Immunofluorescence (IF) analysis

For IF analysis, xrTestes were fixed with 2% paraformaldehyde (Sigma) in PBS for 3 hrs on ice, washed three time with PBS containing 0.2% Tween-20 (PBST), and then immersed successively in 10% and 30% of sucrose (Fisher Scientific) in PBS overnight at 4 °C. The tissues were embedded in OCT compound (Fisher Scientific), frozen and sectioned at a thickness of 10 µm at -20 °C using a cryostat (Leica, CM1800). Sections were placed on Superfrost Microscope glass slides (Thermo Fisher Scientific), air-dried and stored at -80 °C until use. For IF analysis, slides were washed three times with PBS, then incubated with blocking solution (5% normal goat serum in PBST) for 1 hr. The slides were subsequently incubated with primary antibodies in blocking solution for 2 hrs at room temperature followed by washing four times with PBS. They were then incubated with secondary antibodies and 1 µg/ml DAPI in blocking solution for 50 min at room temperature. After washing four times with PBS, the slides were mounted in Vectashield mounting medium (Vector Laboratories) for confocal laser scanning microscopy analysis (Leica, SP5-FLIM inverted).

For IF analyses of fetal testes or some of xrTestes, samples were first fixed in 10% buffered formalin (Fisher Healthcare) with gentle rocking overnight at room temperature. After dehydration, tissues were embedded in paraffin and serially sectioned at a thickness of 4 um using a microtome (Thermo Scientific Microm™ HM325) and placed on Superfrost Microscope glass slides. IF procedure for paraffin sections were similar to that for frozen section with some modification. First, the paraffin sections were de-paraffinized by xylene followed by antigen retrieval by treatment with HistoVT one (Nakalai tesque) for 35 min at 90 °C and then for 15 min at room temperature. After washing with PBS three times, slides were incubated with blocking solution (5% normal donkey serum, 0.2% Tween20 in PBS) for 1 hr at room temperature. The slides were subsequently stained with primary antibodies in blocking solution overnight at 4°C. The slides were then washed 6 times with PBS followed by incubation with secondary antibodies and 1 µg/ml DAPI in blocking solution for 50 min at room temperature. After washing six times with PBS, slides were mounted in Vectashield mounting medium for confocal microscopic analysis.

IF for 5mC was performed using paraffin section as described above with modification. After stained with primary and secondary antibodies other than anti-5mC antibody, slides were treated with 4N HCl in 0.1% Triton X for 10 min at room temperature followed by washing twice briefly with PBS and once for 15 min with PBST before another blocking step. Finally, the slides were incubated with primary antibodies (anti-5mC and other antibodies) followed by secondary antibodies.

### Combined IF and *In Situ* Hybridization (ISH)

ISH on formalin-fixed paraffin-embedded sections was performed using ViewRNA ISH Tissue Assay kit (Thermo Fisher Scientific). Gene-specific probe sets for human *PIWIL4* (VA1-3014459VT), human *ACTB* (VA1-103510VT, positive control) and Bacilus subtilis *dapB* (VF1-11712VT, negative control) were used. The experiments were performed according to the manufacturers’ instruction. Briefly, de-paraffinized sections were treated with pretreatment solution for 10 min at 95 °C, followed by the protease solution for 7 min at 40 °C. Sections were then fixed with 10% buffered formalin for 5 min followed by hybridization with the ViewRNA type 1 probe set for 2 hrs at 40 °C. After vigorous washing with wash buffer, sections were treated with preamplifier probe (25 min at 40°C), Amplifier probe (15 min at 40°C), and the Label Probe 1-AP (15 min at 40°C). Next, sections were incubated with the AP enhancer for 5 min at room temperature followed by treatment of FastRed substrate solution for 1 hr at room temperature. After briefly washing once with PBS, the slides were used for IF studies as described above except without antigen retrieval process. All human testicular tissues were confirmed to be ubiquitously reactive for *ACTB* and negative for *dapB* by confocal imaging (data not shown).

### Fluorescence-activated cell sorting (FACS)

The d5 floating aggregates containing hPGCLCs were incubated with 0.1% Trypsin-EDTA (Gibco) in PBS for 15 min at 37 °C with periodical pipetting. After quenching the reaction by addition of FBS and pipetting, cells were strained through a 70 µm nylon cell strainer (Thermo Fisher Scientific). The AG^+^ cells were sorted by FACSAria Fusion (BD Biosciences) and collected in an Eppendorf tube containing α-MEM.

For the analysis and the sorting of the xrTestes-derived GCs, xrTestes were treated with 0.1% Trypsin-EDTA in PBS for 15 min at 37 °C with periodical pipetting. After quenching the reaction by addition of FBS and pipetting, cell suspension was washed with 0.1% BSA fraction V in PBS and strained through a 70 µm nylon cell strainer. AG^+/-^VT^+/-^PC^+/-^ cells were sorted by FACSAria Fusion and collected in CELLOTION (Amsbio).

### 10x Genomics single-cell RNA-seq library preparation

FACS-sorted *in vitro* cells (hPGCLCs, d81 and d124 xrTestis samples) were collected in CELLOTION. hiPSCs and iMeLCs were collected in StemFit Basic 04 and GK15, respectively without FACS sorting. After centrifuge at 300g for 5 min, cell pellets were resuspended in 0.1% BSA in PBS.

Cells were loaded into Chromium microfluidic chips with Chromium Single cell 3’ v3 chemistry and used to generate single cell gelbead emulsions (GEMs) using Chromium controller (10x Genomics) according to manufacturer’s protocol. GEM-RT was performed in C1000 Touch Thermal Cycler with 96-Deep Well Reaction Module (Bio-Rad). All subsequent cDNA amplification and library construction steps were performed according to manufacturer protocol. Libraries were sequenced using 2 x 150 paired-end sequencing protocol on an Illumina HiSeq 4000 or NovaSeq 6000 instrument.

## QUANTIFICATION AND STATISTICAL ANALYSIS

### Mapping reads of 10x Chromium scRNA-seq and data analysis

Raw data were demultiplexed using mkfastq command in cell Ranger (v3.1.0) to generate Fastq files. Fastq files were trimmed (28bp Cell barcode and UMI Read1, 8bp i7 index, and 91bp Read2) using cutadapt (v.2.6). Trimmed sequence files were mapped on the reference genome for humans (GRCh38-3.0.0). Read counts were obtained from outputs made by CellRanger (v3.1) using edgeR (v3.22.3). Cells with fewer than 1800 genes, and genes with fewer than 10 reads, were filtered out. Genes detected in only five or fewer cells were also omitted from the downstream analyses. Secondary data analyses were performed using R software version (1.2.5019) with the ggplot2 (v3.2.1), gplots (ver.3.0.1.1), qvalue (ver.2.18.0), and sp (ver.1.4-1), TCC (v1.12.1) packages and Excel (Microsoft). All analyses of expression data were performed using log2(TPM+1) values. UHC was performed using hclust function with Euclidian distances and Ward’s method (ward.D2) or complete method (complete). PCA was performed using the prcomp function without scaling. tSNE analysis was performed using Rtsne function and visualized using ggplot2. The RNA velocity analysis was performed using Velocyte ^32^ (http://velocyto.org/) on the cells filtered using the same selection criteria described above.

For identification of DEGs (marker genes) among groups, edgeR function from TCC package was first used to isolate genes with false discovery rate (FDR) < 0.01. The DEGs were then defined as the genes exhibiting more than 2-fold higher expression in one cluster than the remaining clusters. For identification of DEGs in pairwise comparisons, low-expressed genes were first filtered out (mean log2[TPM+1] < 2). Then, Welch-t test and Benjamini-Hochberg method were applied to calculate p-value and FDR, respectively. Among genes with FDR < 0.01, genes with more than 4-fold difference were defined as DEGs, which were plotted over the scatter plot of averaged transcriptome values for cell clusters. For GO analysis, DAVID web tool was used^71^. For evaluating the expression of human infertility genes in our cell clusters, total of 105 genes were retrieved from Cannarella et al., 2020^72^ and Online inheritance in Man (OMIM) database using search categories of “male infertility” and “spermatogenic failure” with subsequent exclusion of genes associated with pituitary, hypothalamus and gonadal somatic cells. Among 105 genes, 45 genes with the average expression value of log2(TPM+1) >1 in at least one cell cluster were selected for analysis.

For comparative study with oogonia-like cells derived from female hiPSCs^18^, gene expression matrix file (log2[RPM+1] values) was retrieved from the article. Given the timing of the culture and AG/VT expression status, we considered that hiPSCs, iMeLCs, hPGCLCs, ag77AG^+^VT^-^, and Ag120 AG^+/-^VT^+^/Ag120AG^-^VT^+^ roughly correspond to our hiPSCs, iMeLCs, hPGCLCs, MLCs and TCs/T1LCs, respectively. We generated the averaged expression values for each cell types in the dataset by Yamashiro et al. using expression values from 2-3 replicates for hiPSCs, iMeLCs, hPGCLCs, ag77AG^+^VT^-^. Ag120 AG^+/-^VT^+^ and Ag120AG^-^VT^+^ samples have no replicate individually, so we lumped these samples and used the averaged value of them for Ag120 AG^+/-^VT^+^ oogonia-like cells. Genes commonly upregulated in T1LCs in this study and ag120 AG^+/-^VT^+^ oogonia-like cells from Yamashiro et.al. were selected using the following criteria. First, among 1268 marker genes for T1LCs defined by DEG analysis as described above (expression 2-fold higher than any other cell cluster in this study, FDR < 0.01), 1177 genes that were also annotated in the dataset by Yamashiro et al. were isolated. Among them, 316 genes showing higher expression over 2-fold in ag120 AG^+/-^VT^+^ oogonia-like cells than any other cell types in the dataset were selected and their expression levels in cell types by Yamashiro et. al. were shown in the heatmap. Genes highly expressed in T1LCs but had weak/no expression in ag120 AG^+/-^VT^+^ oogonia-like cells were selected as follows. First, to enable comparison between two different scRNA-seq platform, RPM values of the data from Yamashiro et al. were adjusted using the polynomial regression curve (y= 0.0908×2-0.0454x+0.1243, least square method) defined using scatter plot comparison between hiPSCs samples from this study and Yamashiro et al. Among 1177 DEGs for T1LCs, 110 genes showing high levels of expression in T1LCs [mean log2(TPM+1) > 4] but had low levels of expression in oogonia-like cells [adjusted log2(RPM+1) < 2] were selected and expression values in cell types in this study and those in Yamashiro et al were shown in the heatmap.

### Profiling of TE expression levels from 10x Chromium scRNA-seq data

To quantify the expression level of TEs at the locus resolution, we first prepared a transcript annotation file (i.e., GTF file) including human genes and TEs, annotated by RepeatMasker (Smit, AFA, Hubley, R & Green, P. *RepeatMasker Open*-4.0. 2013-2015; http://www.repeatmasker.org), as previously described^73^. Read mapping and counting were performed using Cell Ranger with the above annotation file. TE loci detected in ≥ 0.5% of the cells were used in the downstream analyses. The read abundances of total TEs, respective TE families, and TE subfamilies were calculated by summing up the read counts of TE loci. The expression level of TEs was normalized as log2-transformed counts per 10,000 with the pseudocount of 1 (log2 (CP10k + 1)) using Seurat (3.1.4)^74^. Dimension reduction analysis of scRNA-seq data based on TE expression profile was performed according to the Seurat flamework (https://satijalab.org/seurat/). Expression level of TEs was normalized by sctransform^75^. Most variably expressed 100 TE subfamilies were selected by the FindVariableFeatures command. After scaling and whitening data, dimension reduction analysis was performed using Uniform Manifold Approximation and Projection (UMAP)^76^. The first 10 principle components were used in the analysis.

### Epigenetic profiling of TEs

ATAC-Seq data were downloaded from sequence read archive (SRA) using SRA Toolkit (2.10.2) (https://trace.ncbi.nlm.nih.gov/Traces/sra/sra.cgi?view=software). Reads were mapped to the human reference genome (GRCh38-3.0.0) using BWA mem (0.7.17) with the default parameters^77^. Duplicated reads were removed using Picard MarkDuplicates (2.18.6) (https://broadinstitute.github.io/picard/). ATAC-Seq peak was called using MACS2 (2.2.6) with the default parameters^78^. Among called peaks, the top 50,000 peaks were used in the downstream analysis. Fold enrichment value of the overlaps between ATAC-Seq peaks and a specific subfamily of TEs was calculated using an in-house script (calc_enrichment_randomized.py [https://github.com/TheSatoLab/TE_analysis_tools]).

## DATA AND SOFTWARE AVAILABILITY

Accession numbers for the data generated in this study and for the published data used in this study are as follows: the scRNA-seq data in this study (in the process for registration), those of migrating, mitotic and mitotic arrest hFGC (GSE86146)^14^, and neonatal prospermatogonia (GSE124263)^21^, and the RNA-seq data of hPGCLCs and xrOvaries (GSE117101)^18^.

## ACKNOWLEDGMENTS

We thank Drs. Leslie King, Kenneth Zaret and Jeremy Wang for carefully reviewing the manuscript and providing insightful comments. We thank members of Sasaki lab for the discussion of this study and Ms. Karen Makar for her technical and administrative assistance. We thank Dr. Katalin Susztak for letting us use 10X Genomics Chromium Controller. We acknowledge the Comparative Pathology Core at the University of Pennsylvania School of Veterinary Medicine for the preparation of paraffin sections.

This work was supported in part by Open Philanthropy fund from Silicon Valley Foundation to K.S., and NIH R01 HD090007 to BPH. Y.S.H. is supported by National Research Foundation of Korea (NRF-2019R1A6A3A03032063).

## AUTHOR CONTRIBUTIONS

K.S. conceived the project, designed the experiments and wrote the manuscript. B.H. edited the manuscript. Y.S.H. conducted hPGCLC induction, xrTestis culture and analyses. K.S. and Y.S. collected *in vivo* samples and prepared for scRNA-seq. Y.S., Y.S. and K.S. assisted in hPGCLC induction and preparation for xrTestis culture. K.S., S.S., B.H, J.I., H.S., K.S. contributed to the analyses of scRNA-seq.

## DECLARATION OF INTERESTS

The authors declare no competing interests.

## SUPPLEMENTARY TABLES

see separate Excel documents Supplementary Table 1. DEGs between cell clusters in fetal testes

Supplementary Table 2. DEGs between cell clusters *in vitro*

Supplementary Table 3. DEGs between T1 and T1LCs

Supplementary Table 4 Markers for migrating, mitotic and mitotic arrest FGCs defined by Li et.al. 2017

Supplementary Table 5. Comparison of gene expression between T1LCs in this study and ag120AG^+/-^VT^+^ cells by Yamashiro et al. 2018

Supplementary Table 6. Primers used in this study

## Notes

### Competing Interest Statement

The authors have declared no competing interest.

