## Supplementary files for "Reconstitution of prospermatogonial specification *in vitro* from human induced pluripotent stem cells"

**Inventory of Supplementary Information**

**SUPPLEMANTARY FIGURES (Supplementary Fig.1-6)**

Supplementary Fig.1. Identification of cell subsets and their markers in human fetal testes
Supplementary Fig.2. Generation of AGVTPC-knock-in hiPSCs
Supplementary Fig.3. Emergence of GCs with features characteristic of M-prospermatogonia in d77 xrTestes
Supplementary Fig.4. Characterization of GCs derived from hiPSCs
Supplementary Fig.5. DEGs by pairwise comparison between *in vitro* derived cells
Supplementary Fig.6. Key markers of *in vitro* derived cells in comparison with *in vivo* GCs

**LEGENDS TO SUPPLEMENTARY FIGURES**

**SUPPLEMANTARY TABLES, see separate Excel documents**

Supplementary Table 1. DEGs between cell clusters in fetal testes
Supplementary Table 2. DEGs between cell clusters *in vitro*
Supplementary Table 3. DEGs between T1 and T1LCs
Supplementary Table 4. Markers for migrating, mitotic and mitotic arrest FGCs defined by Li et.al. 2017
Supplementary Table 5. Comparison of gene expression between T1LCs in this study and ag120AG$^{+/-}$VT$^{+}$ cells by Yamashiro et al. 2018
Supplementary Table 6. Primers used in this study

#### SUPPLEMENTARY FIGURE LEGENDS

**Supplementary Fig. 1. Identification of cell subsets and their markers in human fetal testes**

**a** Representative section of a fetal testis at 17W3d (Hs31) stained with hematoxylin and eosin (HE). Yellow arrowheads indicate presumptive GCs characterized by large and pale nuclei with prominent nucleoli. Green arrowheads indicate fetal Leydig cells characterized by centrally located round nuclei and abundant eosinophilic cytoplasm. Stroma also contains spindle cells of uncertain origin particularly at regions surrounding seminiferous cords (black arrow). Bar, 50 μm.

**b** Mapping results of three *in vivo* samples analyzed by scRNA-seq.

**c** tSNE plot of all testicular cells in our dataset. Cell cluster identities are determined by feature plots of known markers shown in (**d**). GCs, germ cells; SCs, Sertoli cells; ST, stroma; SMCs, smooth muscle cells; FLCs, fetal Leydig cells; ECs, endothelial cells; Mϕ, macrophages.

**d** Gene expression pattern of markers for indicated cell clusters projected on tSNE plots in (**c**). Both known markers and newly identified markers by DEG analysis in (**e**) are shown.

**e** Heatmap of markers for the indicated cell clusters. Cell clusters identified in (**c**) were used for DEG analysis to identify markers for each cell cluster. Representative markers are shown on the right. The number of DEGs for each cell cluster and the color codes denoting markers for respective cell clusters are shown on the left.

**f** RNA velocity analysis of GCs defined in (**c**). Cluster identifies on this tSNE plot are determined by expression pattens of *TFAP2C* and *PIWIL4*, markers for M and T1, respectively, projected on the tSNE space (left). Lineage trajectory predicted by RNA velocity analysis is shown as arrows over each plot. Dotted lines circumscribe cells representing M (blue) or T1 (purple). A large arrow indicates the overall direction of the individual arrows at the border between M and T1.

**g** tSNE plot from Fig. 1a with cells colored according to their sample origin.

**h** Bar graph showing the relative proportion of M and T1 for each sample.

**i** Scatter plot comparisons of transcriptome data between each testicular sample (Hs26, Hs27, Hs31). Transcriptome comparison using whole GCs (top) or T1 (bottom) are shown. The coefficient of determination ($r^2$) for each pairwise comparison is also indicated.

**j** Scatter plot comparison of the averaged expression values between M and T1 from Hs31 only. Purple, genes higher in T1 (149 genes), cyan, genes higher in M (339 genes) (more than 4-fold difference [flanking diagonal lines], mean log2(TPM+1) > 2, FDR < 0.01). Key DEGs are annotated. $r^2 = 0.876$.

**k** Venn diagram showing the overlap of DEGs (M versus T1 comparison) for data using Hs31 only and data using all 3 *in vivo* samples.

**Supplementary Fig. 2. Generation of AGVTPC-knock-in hiPSCs**

**a** Targeting schemes of human *TFAP2C* (top) and *DDX4* (bottom) loci. *TFAP2C* locus in AG hiPSCs (585B1 1-7) has already been targeted by *2A-EGFP* knock-in construct (Sasaki et al., 2015) and was recombined by Cre over-expression in this study to remove Puromycin selection cassette. The construct for knocking in *2A-tdTomato* into *DDX4* loci was also shown. Black boxes indicate the exons. Arrows indicate PCR screening primers.

**b** Schematic illustration of the human *PIWIL4* locus, and the construct for knocking in *2A-ECFP* into the locus.

**c** Screening by PCR of the homologous recombinants for *TFAP2C-p2A-EGFP* (AG) (top), *DDX4-p2A-tdTomato* (VT) (second), *PIWIL4-p2A-ECFP* (PC) (third), and of random integration of the targeting vector (bottom). Rec., Recombined by Cre; non, non-targeted; P.C., positive control. Clone 13 (red) was selected for AGVTPC hiPSC line (9A13). Yellow arrow, 3kb; Yellow asterisk, 1kb.

**d** PCR genotyping of the genomic DNA isolated from 9A13 hiPSCs for *AG*, *VT* or *PC* alleles and random integration of the targeting vector. Yellow arrow, 3kb. Yellow asterisk, 1kb.

**e** Representative result for the karyotype analysis of 9A13 hiPSCs, showing a normal karyotype (46, XY).

**f** A phase-contrast image of 9A13 hiPSCs. Bar, 200 μm.

**g** IF analysis of 9A13 hiPSCs for DAPI (white), POU5F1 (green, left), SOX2 (red) and NANOG (green, right). Bars, 20 μm.

**h** A phase-contrast image of 9A13 hiPSC-derived iMeLCs. Bar, 200 μm.

**i** BF image and fluorescence images of hPGCLCs derived from 9A13 hiPSCs for AG, VT, PC with a merge. Bar, 200 μm.

**j** FACS analysis of d5 hPGCLCs derived from 9A13 hiPSCs. The percentages of the cells in the indicated gates are shown.

**k** IF images of 9A13 hiPSC-derived d5 hPGCLCs (frozen sections) for TFAP2C (green),

SOX17 (red), POU5F1 (green), NANOG (red) and DAPI (white). Merges are shown on the right. Bars, 20 μm.

**Supplementary Fig. 3. Emergence of GCs with features characteristic of M-prospermatogonia in d77 xrTestes**

**a** Bright field images of xrTestes from day (d) 7 to d120 in the air-liquid interphase (ALI) culture. Bar, 200 μm.

**b** Immunofluorescence (IF) analysis of indicated key proteins (green: GFP and TFAP2C, red: TFAP2C and SOX17) with LAMININ or SOX9 (cyan) (top), and GC markers (red: TFAP2C, cyan: DDX4 and DAZL) in hPGCLC-derived cells (green: GFP) (bottom) in d42 and d77 xrTestes. Bars, 20 μm.

**c** The proportion of TFAP2C$^{+}$DDX4$^{-}$, TFAP2C$^{+}$DDX4$^{+}$, or TFAP2C$^{-}$DDX4$^{+}$ cells in d77 xrTestes as assessed by IF. Percentages with SEM are per section. n, the number of respective cells in four sections.

**d** IF analysis (left) and a box plot showing the fluorescence intensity (with the median and 25th and 75th percentiles, right) of 5-methylcytosine (5mC, red) in hPGCLC-derived cells (green: SOX17) and other somatic cells counterstained with DAPI (white). Bar = 20 μm. hPGCLC-derived cells are outlined by white dotted lines. The statistical significance of the differences between GCs (Germ) and somatic cells (Soma) are evaluated by t-test. ***P < 0.001. n, the number of respective cells counted.

**Supplementary Fig. 4. Characterization of GCs derived from hiPSCs**

**a** Mapping results of six *in vitro* samples analyzed by scRNA-seq in this study.

**b** tSNE plot of *in vitro* derived GCs (same as Fig. 4a) colored based on 4 main clusters (left). Cluster identities are determined based on sample origin and expression of known markers for hiPSCs (*SOX2*), iMeLCs (*SOX2*, *EOMES*), hPGCLCs (*NANOS3*) and prospermatogonia (M and T1) (*NANOS3, DAZL, DND1* and *PIWIL2*) projected on the tSNE plot. Dotted line denotes a cluster (cluster 4) used for subclassification in (**c**).

**c** tSNE plot for subclassification of cluster 4 annotated in (**b**). Three subclusters (4-1, 4-2, 4-3) are newly annotated within this cluster and colored accordingly (left). Cluster identities are determined by markers for M (*KHDC3L*, *POU5F1*, *NANOS3*) and T1 (*TEX15*, *MAGEC2*, *PIWIL4*) defined in Fig. 1a. Newly identified markers (*ASB9, FZD8*) for cluster 4-2, named transitional cells (TCs) are also shown (right).

**d** Expression of *ASB9*, a maker for TCs projected ono the tSNE plot of *in vivo* testicular samples used in Fig. 1a. Dotted line denotes the border between M and T1 cell cluster.
**e** Reclustering of cluster 4 defined in (**b**) projected on tSNE plot used for RNA velocity analysis in Fig. 4b (left). Expression of markers for M, T1 and TCs are projected on the tSNE plot to confirm the identities of each cluster (right).
**f** tSNE plot of *in vitro* derived GCs colored based on the sample origin. Dotted lines circumscribe cell clusters defined in (**b**) and (**c**).
**g** Bar graph showing the proportion of samples in each cell cluster defined in (**b**) and (**c**).
**h** Scatter plot comparison of averaged transcriptome values for two biological replicates of hPGCLCs (hPGCLCs_1, hPGCLCs_2).
**i** UHC of the averaged transcriptome values for cell clusters in this study and neonatal prospermatogonia used in Sohni et al., 2019.
**j** PCA analysis of cells as in (**i**). Color codes for cell clusters are indicated.

**Supplementary Fig. 5. DEGs by pairwise comparison between *in vitro* derived cells**
(left) Scatter plots showing DEGs between indicated samples. The genes upregulated with more than 4-fold differences (FDR < 0.01) in indicated cell clusters are plotted in colors used in Fig. 4a. Key genes, the number of DEGs and coefficient of determination ($r^2$) are denoted. (right) GO analyses of the DEGs. Representative genes in each GO category are shown.

**Supplementary Fig. 6. Key markers of *in vitro* derived cells in comparison with *in vivo* GCs**
Violin plots for the averaged expression values of indicated cell clusters for key DEGs defined in Fig. 4e. As a comparison, averaged expression values of the same genes on *in vivo* testicular GC clusters (M and T1) are shown on the right. Color codes for cell clusters are same as Fig. 1a and 4a.

**SUPPLEMENTARY TABLES, see separate Excel documents**
**Supplementary Table 1.** DEGs between cell clusters in fetal testes

**Supplementary Table 2.** DEGs between cell clusters *in vitro*

**Supplementary Table 3.** DEGs between T1 and T1LCs

**Supplementary Table 4.** Markers for migrating, mitotic and mitotic arrest FGCs defined by Li et.al. 2017,

**Supplementary Table 5.** Comparison of gene expression between T1LCs in this study and ag120AG$^{+/-}$VT$^{+}$ cells by Yamashiro et al. 2018

**Supplementary Table 6.** Primers used in this study

### Supplementary Figure 1

a

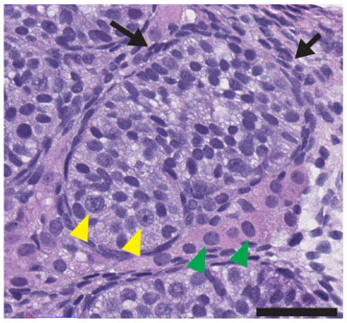

b

| | Total Reads | Mean Reads/cell | Median gene/cell | Detected gene | captured cell | Sequencing Saturation |
|---|---|---|---|---|---|---|
| **Hs26-18W0d** | 177,303,671 | 40,508 | 2,710 | 27,720 | 4,377 | 36.4% |
| **Hs27-18W5d** | 136,780,127 | 22,586 | 1,888 | 26,194 | 6,056 | 27.0% |
| **HS31-17W3d** | 415,576,558 | 69,309 | 2,740 | 26,845 | 5,996 | 60.6% |

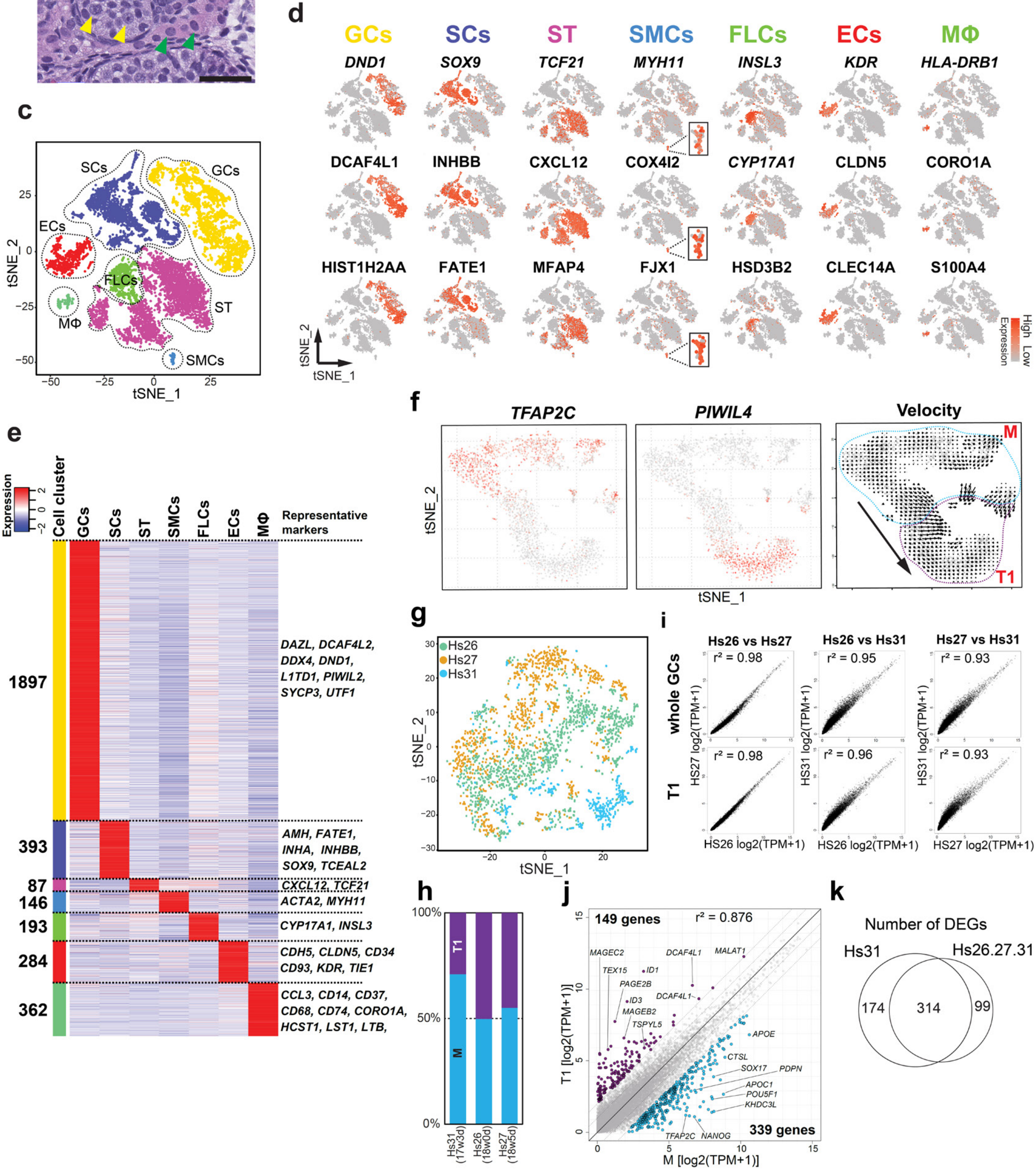

### Supplementary Figure 2

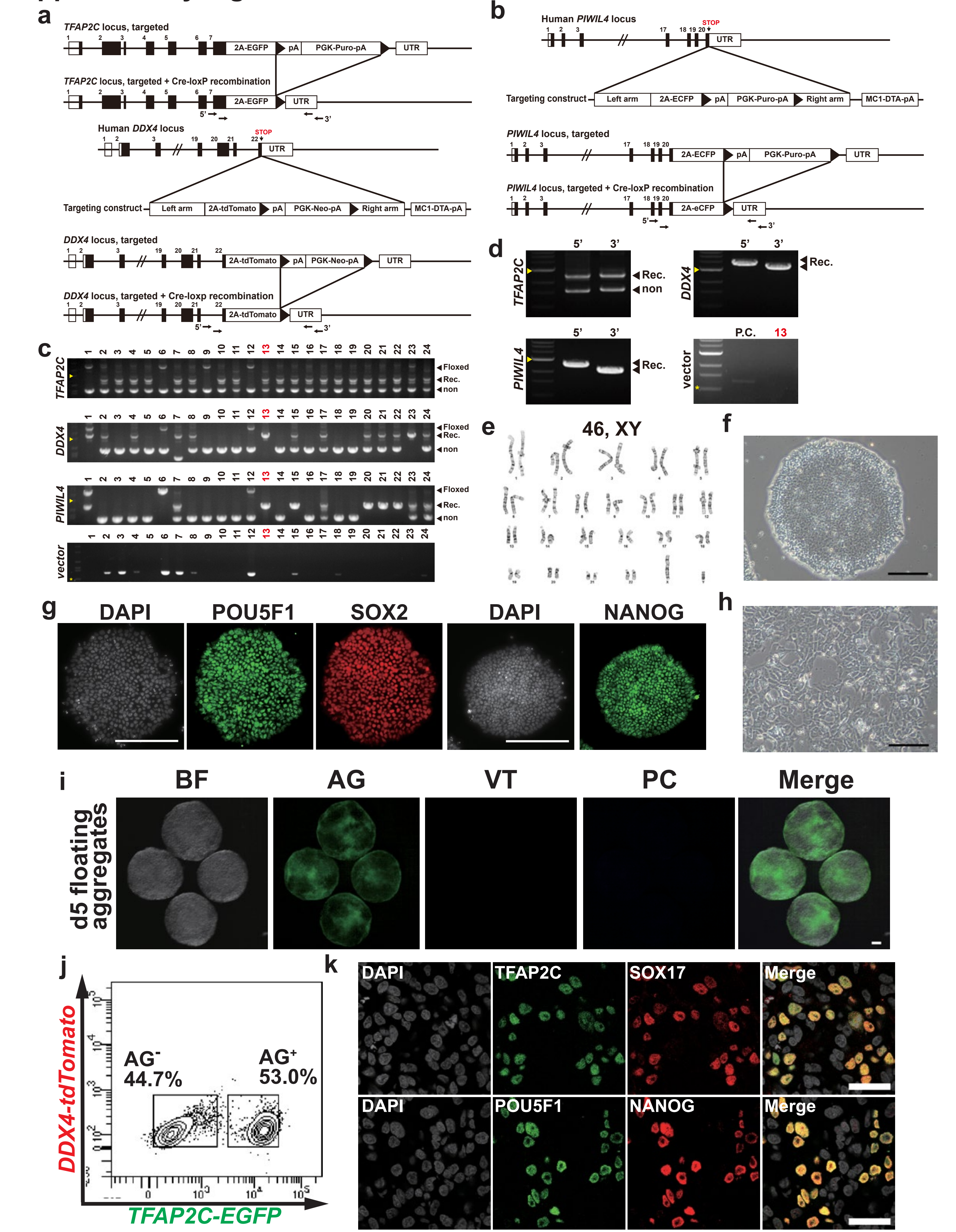

### Supplementary Figure 3

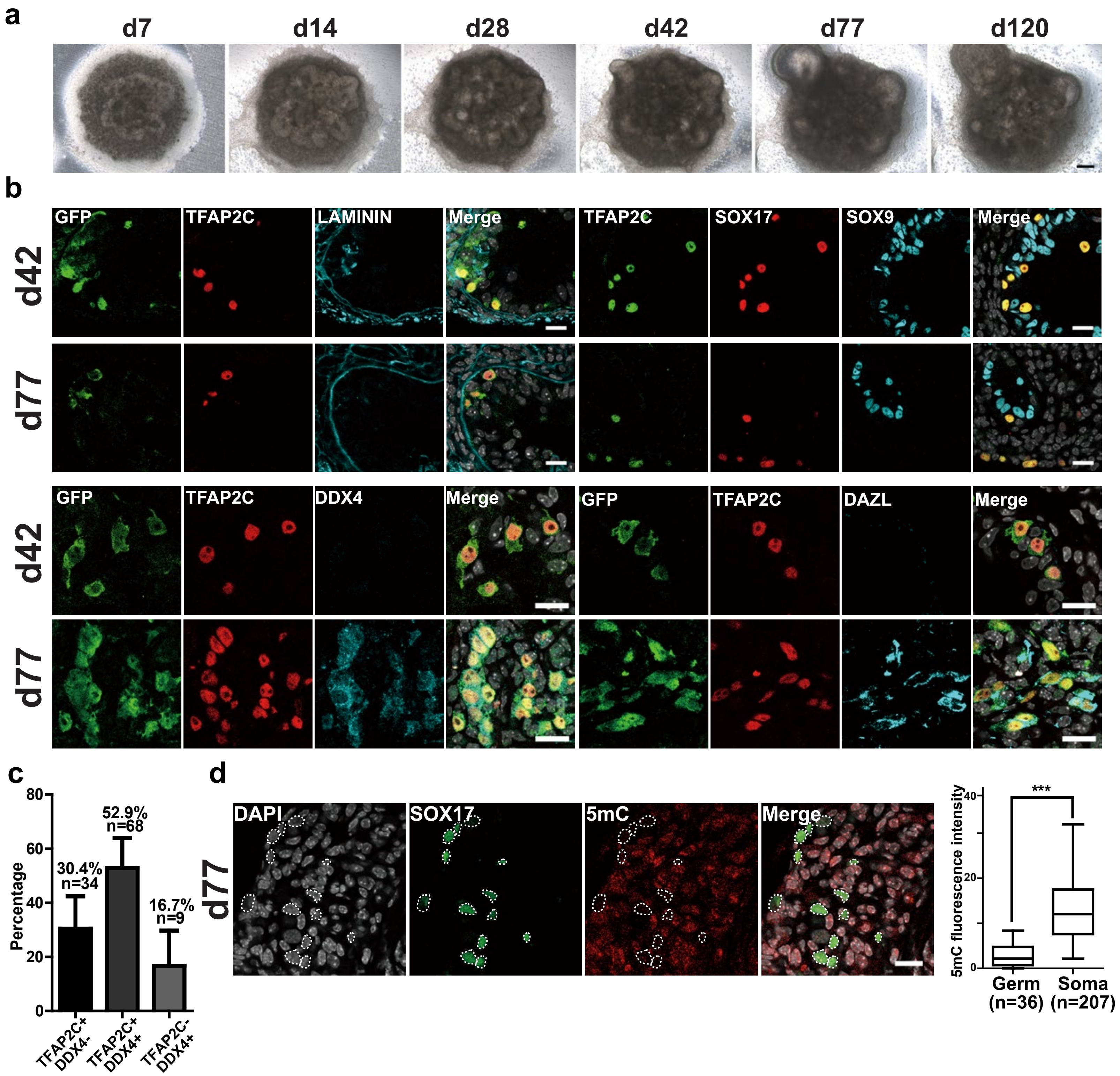

### Supplementary Figure 4

**a**

| | Total Reads | Mean Reads/cell | Median gene/cell | Detected gene | captured cell | Sequencing Saturation |
|---|---|---|---|---|---|---|
| hiPSCs | 85,190,009 | 68,207 | 5,973 | 21,006 | 1,249 | 24.3% |
| iMeLCs | 92,135,422 | 107,761 | 6,843 | 21,255 | 855 | 23.2% |
| hPGCLCs_1 | 107,842,993 | 87,677 | 5,432 | 21,736 | 1,230 | 31.0% |
| hPGCLCs_2 | 34,961,514 | 84,245 | 3,190 | 18,020 | 415 | 48.3% |
| d81_xrTestis | 87,498,123 | 69,943 | 5,010 | 23,123 | 1,251 | 32.6% |
| d124_xrTestis | 93,762,487 | 272,565 | 6,980 | 24,737 | 344 | 63.6% |

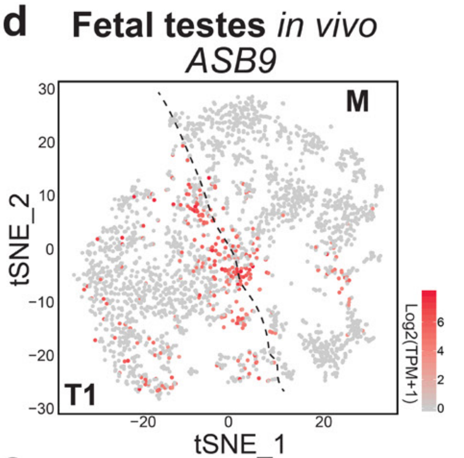


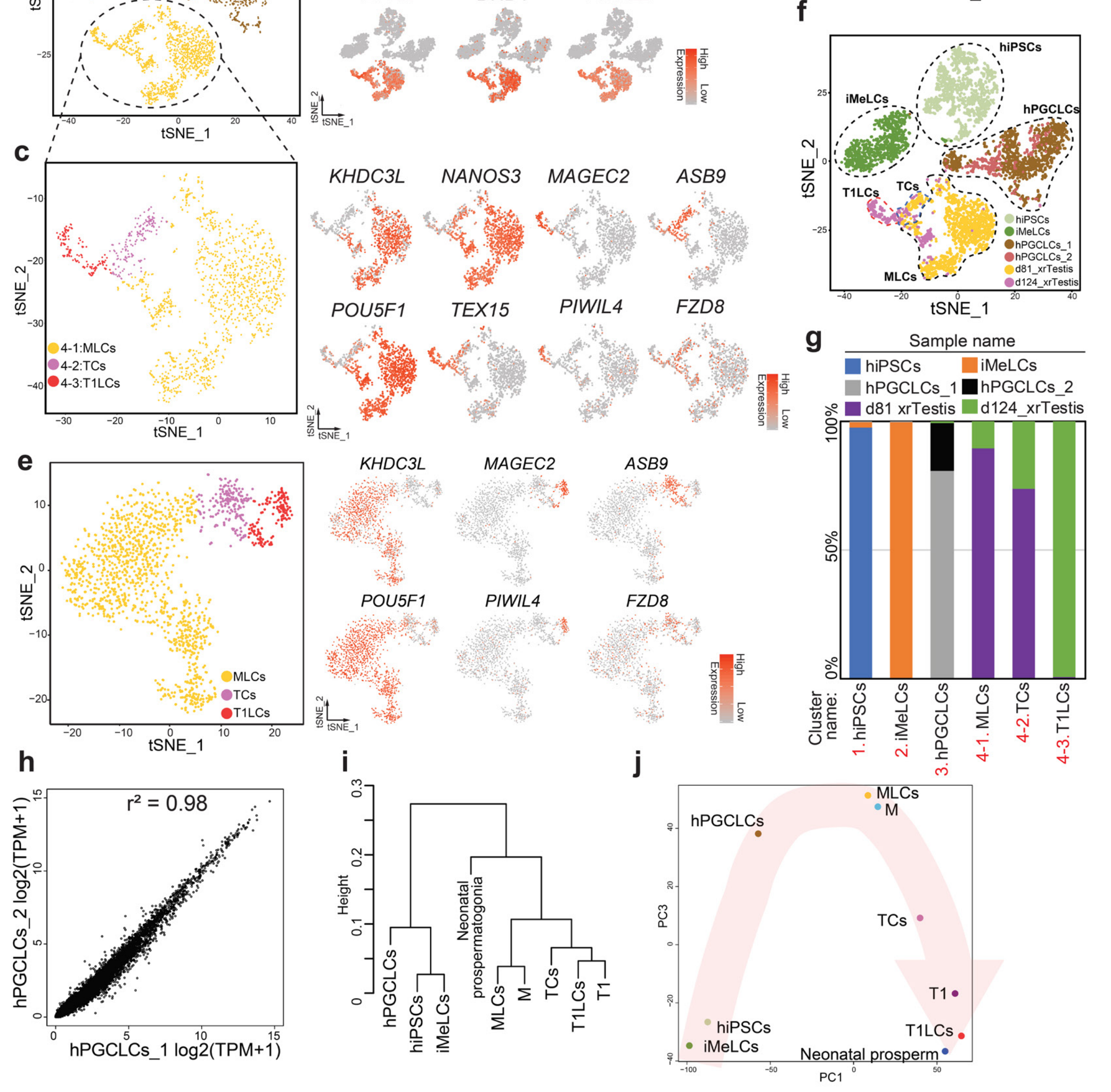

### Supplementary Figure 5

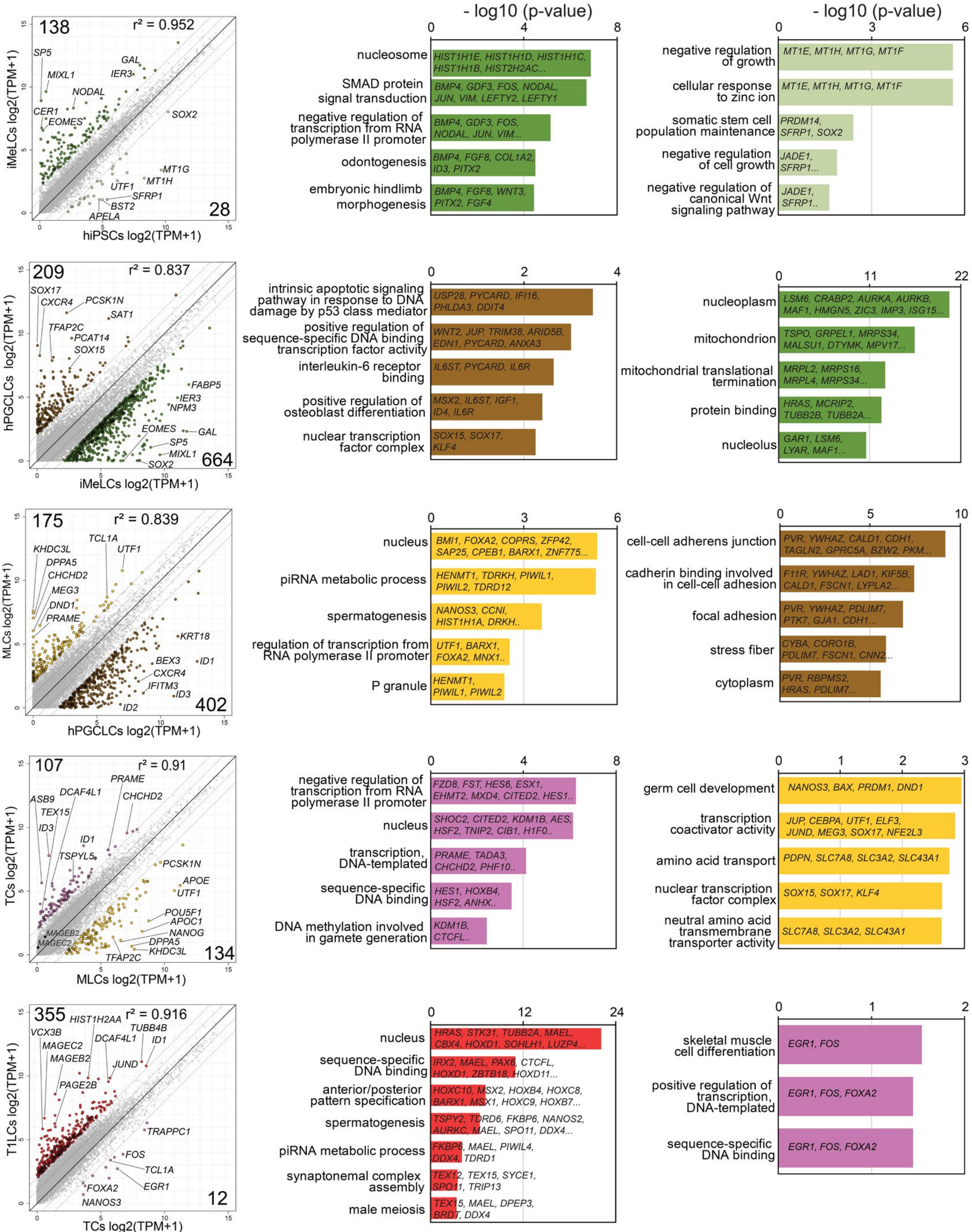

#### Supplementary Figure 6

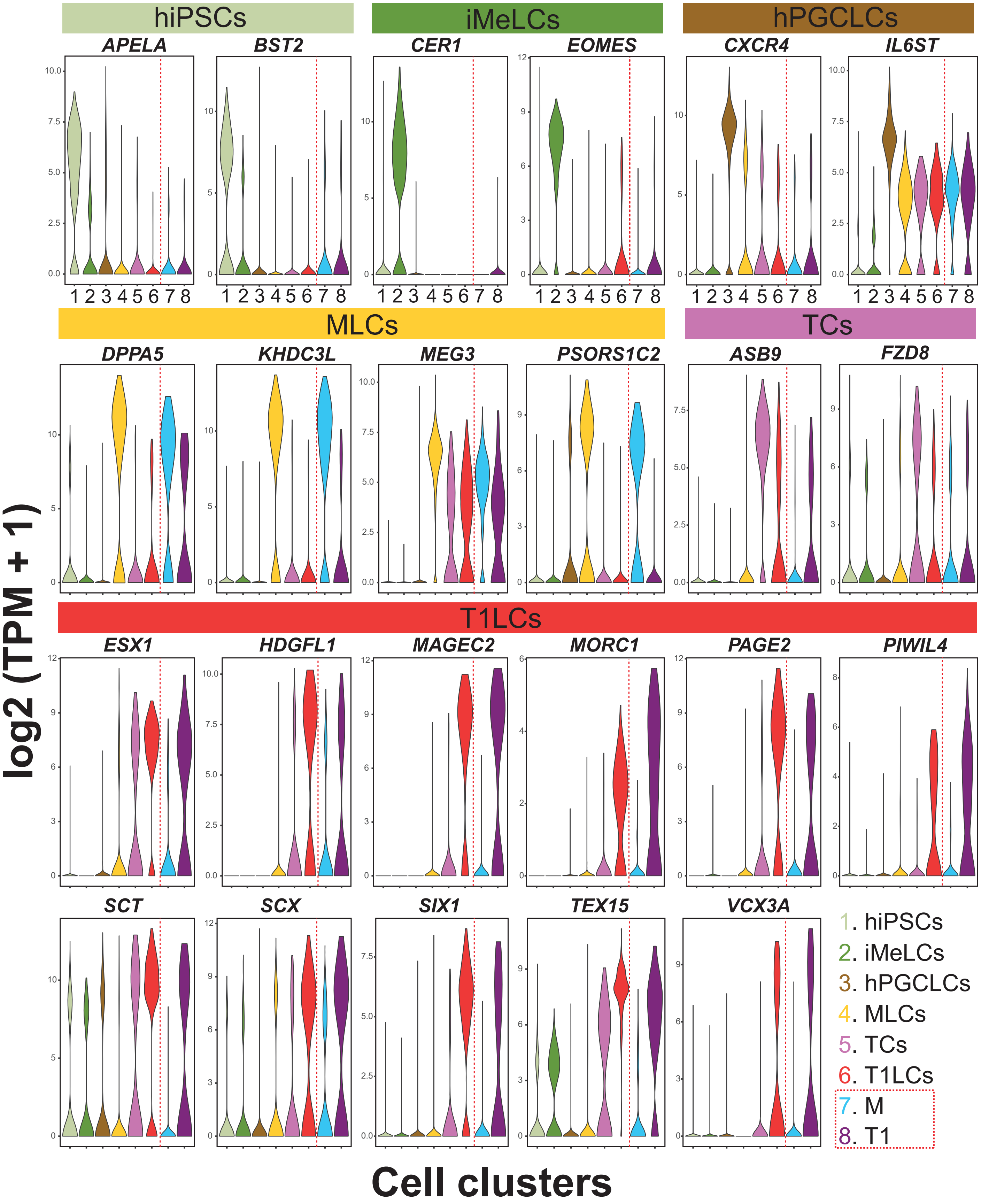
